# The Hedgehog Co-Receptor BOC Differentially Regulates SHH Signaling During Craniofacial Development

**DOI:** 10.1101/2020.02.04.934497

**Authors:** Martha L. Echevarría-Andino, Benjamin L. Allen

## Abstract

The Hedgehog (HH) pathway controls multiple aspects of craniofacial development. HH ligands signal through the canonical receptor PTCH1, and three co-receptors– GAS1, CDON and BOC. Together, these co-receptors are required during embryogenesis to mediate proper HH signaling. Here we investigated the individual and combined contributions of GAS1, CDON and BOC to HH-dependent mammalian craniofacial development. Individual deletion of either *Gas1* or *Cdon* results in variable holoprosencephaly phenotypes, characterized by the failure to divide and form the telencephalon and midfacial structures. In contrast, we find that *Boc* deletion results in facial widening consistent with increased HH pathway activity. Additionally, the deletion of *Boc* in a *Gas1* null background partially rescues the craniofacial defects observed in *Gas1* single mutants; a phenotype that persists over developmental time. This contrasts with HH-dependent phenotypes in other tissues that significantly worsen following combined deletion of *Gas1* and *Boc*. Mechanistically, BOC selectively restricts neural crest-derived mesenchymal proliferation. Together, these data indicate that BOC acts as a multi-functional regulator of HH signaling during craniofacial development, alternately promoting or restraining HH pathway activity in a tissue-specific fashion.

**Summary statement:** Here we identify dual, tissue-specific roles for the Hedgehog co-receptor BOC in both the promotion and antagonism of Hedgehog signaling during craniofacial development.

## Introduction

Hedgehog (HH) signaling regulates the patterning and growth of nearly every tissue in the body (Briscoe and Therond, 2013; McMahon et al., 2003). Aberrant HH pathway activity results in severe birth defects including Holoprosencephaly (HPE), a defect characterized by the failure of the division of the embryonic forebrain into two cerebral hemispheres (Muenke and Beachy, 2000). HPE is one of the most common birth defects in humans, estimated to affect as many as 1 in 250 embryos (Hong and Krauss, 2018). The clinical manifestations of HPE are highly heterogeneous, consisting of a wide phenotypic spectrum of defects (Schachter and Krauss, 2008). Notably, 80% or more of HPE cases will display facial defects in addition to forebrain malformations (Schachter and Krauss, 2008).

Multiple mutations associated with developmental signaling pathways such as HH, have been identified in human HPE patients (Roessler and Muenke, 2010). Specifically, mutations in *Sonic Hedgehog* (*SHH)* account for 6%-8% of sporadic HPE (Roessler et al., 2009). During craniofacial development *Shh* regulates the establishment of forebrain identity, and patterns the face primordia (Schachter and Krauss, 2008). Moreover, disruption of *Shh* in mice results in abnormal dorsoventral patterning in the neural tube, defective axial skeleton formation and alobar HPE (Chiang et al., 1996).

SHH ligands signal through the twelve-pass transmembrane receptor Patched (PTCH1), (Marigo et al., 1996). However, SHH also binds three co-receptors, growth arrest specific 1 (GAS1), CAM- related/downregulated by oncogenes (CDON) and brother of CDON (BOC) (Allen et al., 2011; Allen et al., 2007; Beachy et al., 2010; Izzi et al., 2011; Lee et al., 2001; McLellan et al., 2008; Tenzen et al., 2006; Yao et al., 2006; Zhang et al., 2011; Zhang et al., 2006). CDON and BOC are structurally similar members of the immunoglobulin superfamily that are conserved from *Drosophila* to mammals (Beachy et al., 2010; Kang et al., 1997; Kang et al., 2002; Lum et al., 2003). GAS1 is a vertebrate-specific, GPI-anchored protein with structural resemblance to GDNF receptors (Cabrera et al., 2006). In the absence of SHH ligand, PTCH1 inhibits the activity of the GPCR-like protein Smoothened (SMO). SHH ligand binding to PTCH1 and GAS1, CDON or BOC releases SMO inhibition leading to a signal transduction cascade that results in modulation of the GLI family of transcriptional effectors (Hui and Angers, 2011). Together, GAS1, CDON and BOC are required for HH signal transduction during embryogenesis (Allen et al., 2011; Allen et al., 2007; Cole and Krauss, 2003; Izzi et al., 2011; Martinelli and Fan, 2007; Tenzen et al., 2006; Zhang et al., 2011; Zhang et al., 2006)

Similar to *Shh* mutants, simultaneous genetic removal of *Gas1*, *Cdon* and *Boc* results in alobar HPE (Allen et al., 2011). Further, multiple mutations in these HH co-receptors have been identified in human HPE patients (Bae et al., 2011; Hong et al., 2017; Ribeiro et al., 2010), suggesting that these proteins play key roles in craniofacial development. This is supported by multiple studies in mice demonstrating a role for these genes during HH-dependent craniofacial development (Cole and Krauss, 2003; Seppala et al., 2007; Seppala et al., 2014; Zhang et al., 2011; Zhang et al., 2006). *Gas1* and *Cdon* single mutants display microforms of HPE, in which the severity of the phenotype is dependent on the genetic background of the mouse model (Allen et al., 2007; Cole and Krauss, 2003; Seppala et al., 2007; Zhang et al., 2006). In contrast, in mixed genetic backgrounds *Boc* deletion does not result in any HPE phenotypes, although these animals do display defects in SHH-dependent commissural axon guidance (Okada et al., 2006; Seppala et al., 2014; Zhang et al., 2011). More recently, *Boc* has been demonstrated to function as silent HPE modifier gene that, in the context of other HPE mutations, can modify the severity of the HPE phenotype (Hong and Krauss, 2018). It has been proposed that modifier genes like *Boc* contribute to the phenotypic differences observed in different genetic backgrounds.

GAS1, CDON and BOC have generally been described as positive regulators of the HH signaling pathway. However, in certain contexts these co-receptors can act to restrain HH signaling. For example, *Gas1* can antagonize HH signaling in presomitic mesoderm explants (Lee et al., 2001), and restricts HH signaling during tooth development in mice (Cobourne et al., 2004; Ohazama et al., 2009). Similarly, *Cdon* negatively regulates HH pathway function in the optic vesicle of zebrafish and chicken embryos (Cardozo et al., 2014), while *Boc* antagonizes HH signaling in the zebrafish lower jaw (Bergeron et al., 2011). It remains unclear how these co-receptors differentially regulate HH signaling in these different contexts.

Here we investigated the contributions of GAS1, CDON and BOC to HH-dependent mammalian craniofacial development. Specifically, we examined the individual and combined deletion of different HH co-receptors on a congenic C57BL/6J background. Surprisingly, we found that *Boc* mutants display facial widening, consistent with HH increased activity. Additionally, deletion of *Boc* in a *Gas1* null background partially ameliorates the craniofacial defects observed in *Gas1* single mutants, while other HH-dependent phenotypes in these mutants are significantly worsened. Interestingly, the rescue of the craniofacial defects in *Gas1;Boc* mutants persists over developmental time, and is restricted to the nasal bone and the nasal capsule. Finally we provide evidence that BOC selectively restricts neural crest-derived mesenchymal proliferation. Together, our data indicate that BOC acts as a multi-functional regulator of HH signaling during craniofacial development, alternately promoting or restraining HH pathway activity in a tissue-specific fashion.

## Results

To define the expression of the HH pathway co-receptors *Gas1*, *Cdon* and *Boc* during early craniofacial development, we utilized *lacZ* (*Gas1* and *Cdon*) and *Alkaline phosphatase* (*AP*; *Boc*) reporter alleles (Fig. 1) (Cole and Krauss, 2003; Martinelli and Fan, 2007; Zhang et al., 2011). At E8.5 *Gas1*, *Cdon* and *Boc* are primarily expressed in the cranial neural folds, the somites and the neural tube (Fig. 1A-D). During this stage *Cdon* is the only co-receptor expressed in the prechordal plate (PCP), a major signaling center during craniofacial development that secrets SHH ligand, which patterns the ventral telencephalon (Fig. 1C) (Cordero et al., 2004; Rubenstein and Beachy, 1998; Zhang et al., 2006). As development progresses, these expression patterns are maintained in the somites and neural tube, and expand to additional structures. At E9.5 the HH co-receptors are all expressed in the frontonasal prominence (FNP), maxillary process (MXP) and mandibular process (MP; Fig. 1E-H). Differences in *Gas1*, *Cdon* and *Boc* expression in craniofacial structures are revealed by analysis of E10.5 embryos (Fig. 1I-T).

**Figure 1.**
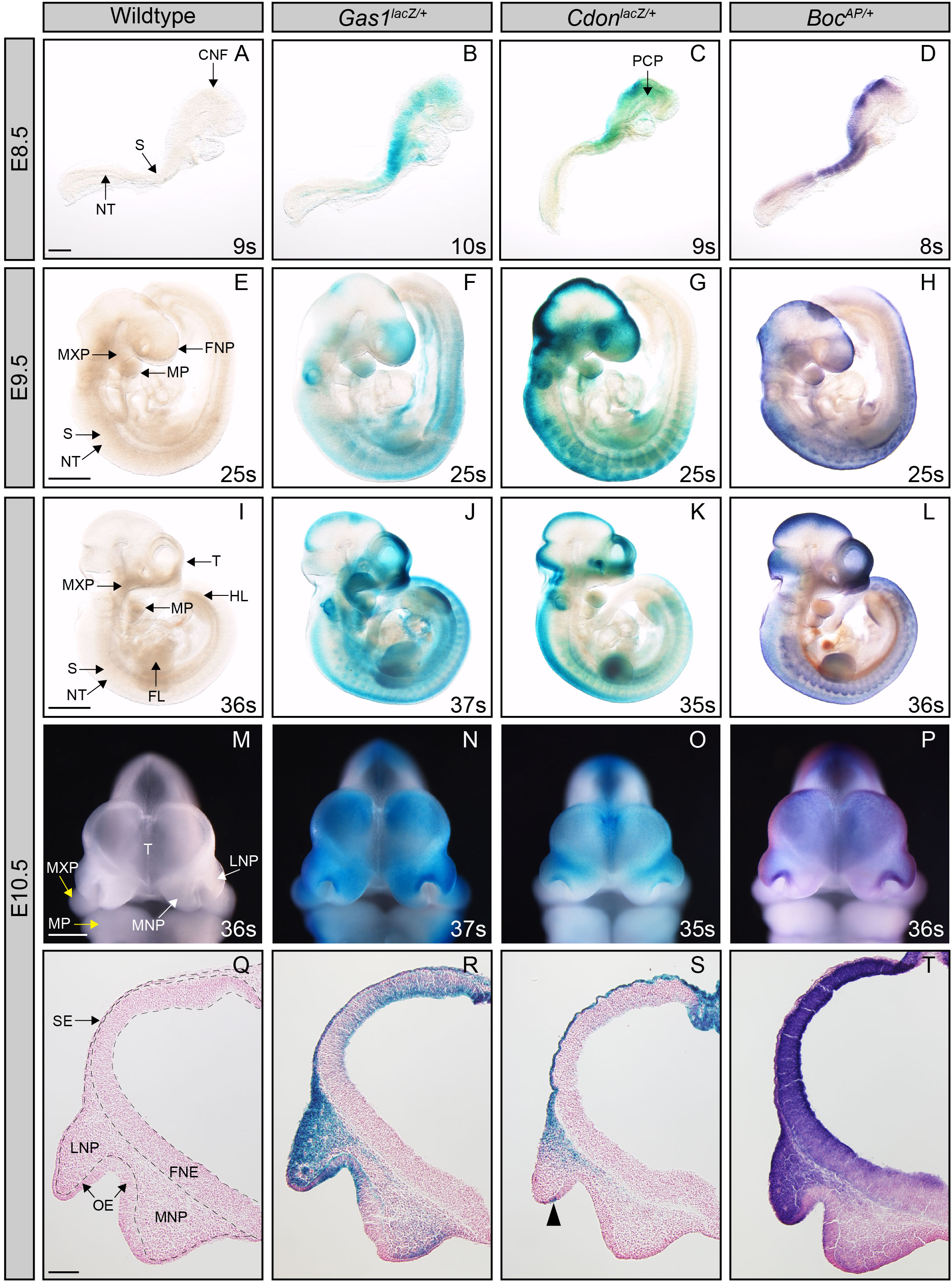
The HH co-receptors *Gas1*, *Cdon* and *Boc* are expressed throughout early craniofacial development. Analysis of HH co-receptor expression using *lacZ* (*Gas1*, *Cdon*) and *hPLAP* (*Boc*) reporter alleles (A-T). Whole mount X-Gal and Alkaline Phosphatase staining of E8.5 (A-D), E9.5 (E-H), and E10.5 (I-L), wildtype (A, E, I, M, Q), *Gas1^lacZ/+^* (B, F, J, N, R), *Cdon^lacZ/+^* (C, G, K, O, S), and *Boc^AP/+^* (D, H, L, P, T) embryos is shown. Somite number (s) is indicated in the lower right corner of each panel. Frontal view of craniofacial structures of E10.5 embryos (M-P). White arrows denote LNP and MNP and yellow arrows denote MXP and MP (M). Coronal sections of E10.5 telencephala (Q-T); arrowhead denotes a subset of cells expressing *Cdon* in the olfactory epithelium. Scale bars, (A-P) 500 µm and (Q-T) 200 µm. Abbreviations: cranial neural fold (CNF), somites (S), neural tube (NT), pre-chordal plate (PCP), frontonasal prominence (FNP), maxillary process (MXP), mandibular process (MP), telencephalon (T), forelimb (FL), hindlimb (HL), medial nasal process (MNP), lateral nasal process (LNP), surface ectoderm (SE), neuroepithelium (NE) and olfactory epithelium (OE).

En face views of whole-mount stained E10.5 embryos (Fig. 1M-P) demonstrate broad expression of *Gas1, Cdon* and *Boc* in the telencephalon. X-Gal and AP staining in coronal sections of E10.5 embryos reveals that all three co-receptors are present in the surface ectoderm and in the neuroepithelium (NE) of the telencephalon in a dorso-ventral gradient (Fig. 1 Q-T; Fig. S1A-D). Notably, the ventral extent of *Cdon* expression in the NE is restricted compared to *Gas1* and *Boc*. Similarly, *Gas1* and *Boc* display broad expression in the olfactory epithelium (OE), while *Cdon* expression is limited to a subset of cells in the medial OE of the LNP (see black arrowhead in Fig. 1S, Fig. S1F).

At E10.5, *Gas1* is the only co-receptor expressed in the MP and in the MXP (Fig. 1J, N). Further differences in the expression of the HH co-receptors are detected in the medial nasal and lateral nasal processes (MNP and LNP). All three co-receptors are expressed in the LNP (Fig. 1Q-T). However, *Gas1* and *Boc* are expressed throughout the LNP mesenchyme, while *Cdon* expression is restricted to the most dorsal aspect of the LNP mesenchyme (Fig. 1S, Fig. S1F). In the MNP, *Gas1* and *Boc* are broadly expressed at lower levels in the mesenchyme; in contrast, *Cdon* is only expressed in mesenchymal cells that are proximal to the NE (Fig. 1S, Fig. S1F). The expression patterns of *Gas1, Cdon* and *Boc* in craniofacial structures are consistent with their negative transcriptional regulation by the HH signaling pathway (Allen et al., 2007; Tenzen et al., 2006). In addition to differences in expression of the HH co-receptors in craniofacial structures, their expression in other HH-responsive tissues such as the forelimb bud (Fig. S1H– K), and the neural tube (Fig. S1L-O) is also not identical. Overall, we noted that the expression domain of *Boc* in the NE of the telencephalon and in the neural tube extends closer to the SHH ligand source in both tissues. These data raise the question of whether these co-receptors, and BOC in particular, may differentially contribute to HH-dependent craniofacial development.

To address the individual contributions of *Gas1*, *Cdon*, and *Boc* to craniofacial development, we examined single mutant embryos at mid-gestation on a congenic C57BL/6J background (Fig. 2). At E10.5 *Gas1^-/-^* and *Cdon^-/-^* embryos display a spectrum of HPE phenotypes that range from proper telencephalic vesicle (TV) division with normal MNP separation, to no TV division with no MNP separation (Fig. 2A-D, E-H; Fig. S2). Most of these mutants exhibit incomplete TV division (76% of *Gas1*^-/-^ embryos, and 50% of *Cdon^-/-^* embryos), while a smaller portion (12% and 17%, respectively) of these mutants fails to divide the TV (Fig. 2M). *Gas1^-/-^* and *Cdon^-/-^* embryos predominantly show either incomplete MNP separation (47% of *Gas1*^-/-^ embryos, and 33% of *Cdon^-/-^* embryos) or no MNP separation (29% and 42%, respectively; Fig. 2N). Notably, a minority of *Gas1* and *Cdon* mutants have more mild phenotypes that are characterized by normal TV division (Fig. 2M) and either normal or reduced MNP separation (Fig. 2N). In contrast, *Boc^-/-^* embryos do not manifest any gross craniofacial defects (Fig. 2I-L), with 100% of embryos displaying normal TV division and normal MNP separation (Fig. 2M-N). Together, these data indicate that even on a congenic C57BL/6J genetic background there remains a spectrum of HPE phenotypes observed in *Gas1* and *Cdon* mutants. Strikingly, and despite the broad expression of *Boc* in multiple HH-responsive cell types in the developing forebrain (Fig. 1), we do not observe any HPE phenotypes in *Boc* mutants maintained on a C57BL/6J background.

**Figure 2.**
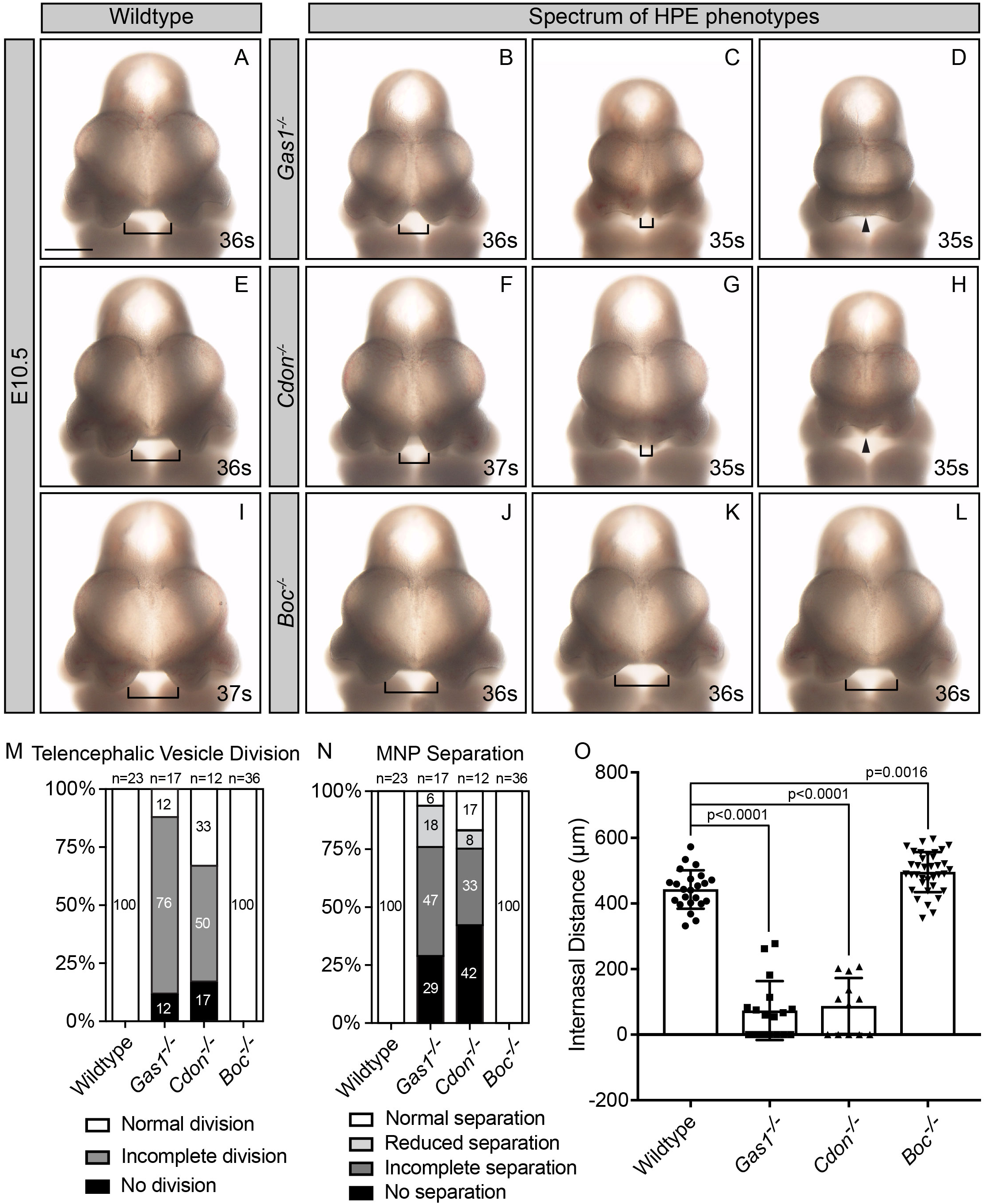
Loss of *Boc* results in midface widening at E10.5 on a congenic C57BL/6J background. En face view of E10.5 mouse embryos (A-L). Somite number (s) is indicated in the lower right corner of each panel. Brackets indicate internasal distance. Black triangles denote fusion of the MNP. E10.5 wildtype (A,E,I), *Gas1^-/-^* (B-D), *Cdon^-/-^* (F-H), and *Boc^-/^*^-^ (J-L) embryos. Note that *Gas1* and *Cdon* mutants display a range of craniofacial defects (increasing in severity from left to right), while *Boc* mutants do not display any gross morphological changes. Scale bar (A), 500 µm. Telencephalic vesicle (TV) division frequency in E10.5 wildtype, *Gas1^-/-^*, *Cdon^-/-^*, and *Boc^-/-^* embryos (M). TV division was classified according to the following categories: normal division, incomplete division and no division (see Fig. S2A-C for a representative example of each category). Medial nasal process (MNP) separation frequency in E10.5 wildtype, *Gas1^-/-^*, *Cdon^-/-^*, and *Boc^-/-^* embryos (N). MNP separation in each embryo was classified according to the following categories: normal separation, reduced separation, incomplete separation and no separation (see Fig. S2D-G for a representative example of each category). Internasal distance quantitation in wildtype (n= 23), *Gas1^-/-^* (n=17), *Cdon^-/-^* (n=12), *Boc^-/-^* (n=36) embryos (O; in µm). Data are represented as the mean ± standard deviation. P-values were determined by a two-tailed Student’s *t*-test.

To further characterize the spectrum of HPE phenotypes, we quantified the internasal distance in E10.5 embryos. Consistent with our initial assessment, this quantitation revealed significant reductions in the internasal distance in both *Gas1* and *Cdon* mutant embryos (Fig. 2O). Surprisingly, this quantitation also revealed an unexpected subtle, but significant increase in the internasal distance in *Boc* mutant embryos compared to wildtype embryos (443μm in wildtype embryos and 496μm in *Boc^-/-^* embryos; Fig. 2O). These data suggest potentially opposing roles for *Gas1* and *Cdon* compared to *Boc* during mammalian craniofacial development. One explanation for these counterintuitive results is that the increased internasal distance in *Boc* embryos was due to an overall increase in embryo size. Therefore, we measured the crown-rump length (CRL) in E10.5 wildtype and mutant embryos (Fig. S3A-E). While *Gas1* mutants are significantly smaller than their wildtype littermates, both *Cdon* and *Boc* mutant embryos have similar CRL as wildtype embryos (Fig. S3F). These data support the notion that the MNP widening observed in *Boc* mutants at E10.5 reflects differences in the contribution of this HH co-receptor to craniofacial development. Interestingly, widening or duplication of midfacial tissues is associated with increased levels of HH signaling (Brugmann et al., 2010; Hu and Helms, 1999).

To determine if the variable craniofacial defects observed in these HH co-receptor mutant embryos correlates with HH pathway activity, we performed *in situ* hybridization for *Gli1*, a general and direct transcriptional target of HH signaling (Dai et al., 1999). *Gli1* is expressed in multiple craniofacial structures, including the MNP, MXP and MP (Fig. S4A). *Gas1^-/-^* and *Cdon^-/-^* embryos with less severe HPE phenotypes maintain *Gli1* expression in the MNP, but embryos with increasingly severe HPE phenotypes display a loss of *Gli1* expression in the MNP (Fig. S4D-F, G-I). Consistent with the midfacial widening observed in *Boc^-/-^* embryos, *Gli1* expression is maintained in the MNP across all *Boc* mutant embryos (Fig. S4J-L). Taken together, these data demonstrate that HPE severity in *Gas1* and *Cdon* mutant embryos correlates with *Gli1* loss in the MNP, and confirms that *Boc* mutants do not display any reduced HH pathway activity during craniofacial development.

While previous studies suggested that combinatorial deletion of *Gas1*, *Cdon*, or *Boc* results in more severe HPE phenotypes (Allen et al., 2011; Allen et al., 2007; Seppala et al., 2014; Zhang et al., 2011), work in zebrafish suggested a potential negative role for *Boc* in lower jaw development (Bergeron et al., 2011). Furthermore, a *Boc* missense variant associated with increased HH pathway activity has been recently identified in human HPE patients (Hong et al., 2017). The midface widening that we observed in *Boc^-/-^* embryos (Fig. 2O) is consistent with a role for *Boc* as a potential HH antagonist during craniofacial development. To explore this possibility, we deleted *Boc* in combination with *Gas1* deletion on a congenic C57BL/6J background.

Analysis of E10.5 *Gas1^-/-^;Boc^-/-^* embryos revealed a spectrum of HPE phenotypes, as observed in *Gas1^-/-^* embryos (Fig. S4M-O). Importantly, the HPE phenotypes observed in *Gas1;Boc* double mutants are less severe than those observed in *Gas1* single mutants (cf. Fig. 3B and 3D). Specifically, we observed an increase in the percentage of *Gas1;Boc* double mutants with normal TV division compared to *Gas1* single mutants (31% vs. 12%, respectively; Fig. 3E). Further, we found that 50% of *Gas1;Boc* double mutants display MNP separation compared to 24% of *Gas1* mutants (Fig. 3F). To investigate whether this rescue was due to increased overall embryo size, we measured the CRL of *Gas1^-/-^;Boc^-/-^* embryos (Fig. S5A-E). We find that *Gas1^-/-^;Boc^-/-^* embryos tend to be smaller than *Gas1^-/-^* embryos (Fig. 5F); while not statistically significant, these data rule out increased embryo size as an explanation for the rescue of the HPE phenotypes. Overall, these data suggest that *Boc* deletion in a *Gas1* mutant background partially rescues TV and MNP separation in E10.5 embryos.

**Figure 3.**
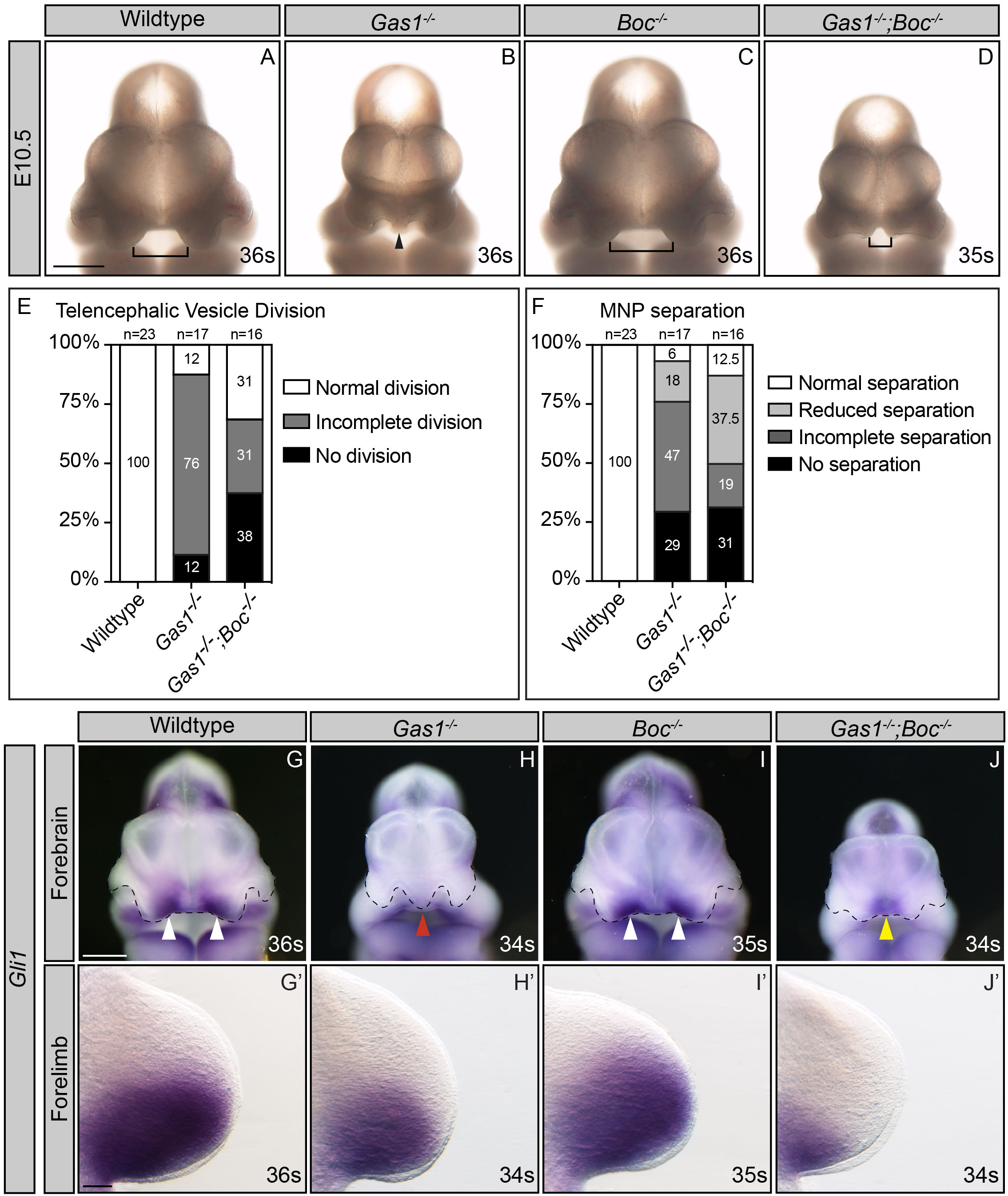
Tissue-specific rescue of HH signaling in E10.5 *Gas1;Boc* double mutant embryos. En face view of E10.5 embryos (A-D). Somite number (s) is indicated in the lower right corner of each panel. Brackets indicate internasal distance. Black triangles denote fusion of the medial nasal process. E10.5 wildtype (A), *Gas1^-/-^* (B), *Boc^-/-^* (C), and *Gas1^-/-^;Boc^-/^*^-^ (D) embryos. Telencephalic vesicle (TV) division frequency in E10.5 wildtype, *Gas1^-/-^*, *Boc^-/-^*, and *Gas1^-/-^*;*Boc^-/-^* embryos (E). TV division was classified according to the following categories: normal division, incomplete division and no division (see Fig. S2A-C for representative examples of each category). Medial nasal process (MNP) separation frequency in E10.5 wildtype, *Gas1^-/-^*, *Boc^-/-^*, and *Gas1^-/-^*;*Boc^-/-^* embryos (F). MNP separation in each embryo was classified according to the following categories: normal separation, reduced separation, incomplete separation and no separation (see Fig. S2D-G for representative examples of each category). *In situ* hybridization detection of *Gli1* expression in E10.5 forebrains (G-J) and their corresponding forelimbs (G’-J’). White arrowheads denote *Gli1* expression in the MNP (G,I); red arrowhead indicates the absence of *Gli1* expression (H); yellow arrowhead marks partial rescue of *Gli1* expression (J). En face view of E10.5 forebrains and dorsal view of E10.5 forelimbs in wildtype (G, G’), *Gas1^-/-^* (H,H’), *Boc^-/-^* (I,I’), and *Gas1^-/-^;Boc^-/-^* (J,J’) embryos. Somite number (s) is indicated in the lower right corner of each panel. Black dotted lines outline nasal processes. Note that *Gli1* is differentially regulated in the MNP and forelimb of *Gas1;Boc* mutants. Scale bar in (A) and (G), 500µm; (G’), 100µm.

To determine if the phenotypes observed in *Gas1;Boc* mutants correlate with changes in HH pathway activity, we performed *in situ* hybridization for the direct HH transcriptional target *Gli1* in E10.5 wildtype, *Gas1^-/-^, Boc^-/-^*, and *Gas^-/-^;Boc^-/-^* embryos (Fig 3G-J). *Gas1^-/-^;Boc^-/-^* embryos that display increased MNP separation also display increased *Gli1* expression in the MNP (Fig 3J), consistent with the notion that *Boc* antagonizes HH pathway activity during craniofacial development. We also examined *Gli1* expression in the forelimb bud from these same embryos (Fig. 3G’-J’). Notably, we detected decreased *Gli1* expression in *Gas1;Boc* mutants compared to wildtype and *Gas1* mutant embryos (cf. Fig. 3G’, H’, J’). Together these data suggest that loss of *Boc* partially and selectively rescues HPE phenotypes observed in *Gas1* mutant embryos, through increased HH pathway activity.

To examine the consequences of *Boc* deletion on additional targets of the HH pathway, and to begin to dissect possible tissue-specific contributions to craniofacial development, we investigated HH-dependent neural patterning in both the developing forebrain and spinal cord (Fig. 4). Specifically, we used whole mount immunofluorescence to analyze the expression of NKX2.1, a direct HH transcriptional target in the ventral telencephalon (Pabst et al., 2000) (Fig. 4E-H,M). In E10.5 *Gas1^-/-^* embryos the expression domain of NKX2.1 is significantly reduced (Fig. 4F), while the NKX2.1+ domain in *Boc^-/-^* embryos is unchanged compared to wildtype embryos (cf. Fig. 4E,G). Notably, compared to *Gas1^-/-^* embryos (Fig. 4F), *Gas1^-/-^;Boc^-/-^* embryos maintain a similar NKX2.1+ domain (Fig. 4H). Quantitation confirms that the NKX2.1+ area is not significantly altered in *Gas1^-/-^;Boc^-/-^* embryos compared to *Gas1^-/-^* embryos (Fig. 4M). We also confirmed that NKX2.1 is not significantly different in *Boc^-/-^* embryos (Fig. 4M). Together, these data suggest that, despite its broad expression in the forebrain neuroepithelium (Fig. 1T), *Boc* does not positively contribute to HH-dependent patterning in this tissue. These data do raise the question of whether *Boc* can regulate HH signaling in the developing telencephalon, or whether it may be playing an antagonistic role. To address these possibilities, we used chicken *in ovo* telencephalon electroporations to assess *Boc* function during HH-dependent neural patterning in the forebrain (Fig. S6). Expression of GFP (pCIG, empty vector) in the chicken telencephalon does not affect NKX2.1 expression (Fig. S6A-D). In contrast, expression of *SmoM2* (a constitutively active form of SMO) (Xie et al., 1998), which drives high levels of HH pathway activity, induces ectopic NKX2.1 expression (Fig. S6E-H). Similarly, expression of *Boc* also induces ectopic NKX2.1 expression (Fig. S6I-L). These data demonstrate that *Boc* can promote HH-dependent patterning in the developing chicken forebrain, and suggests that *Boc* does not play an antagonistic role in the forebrain neuroepithelium.

**Figure 4.**
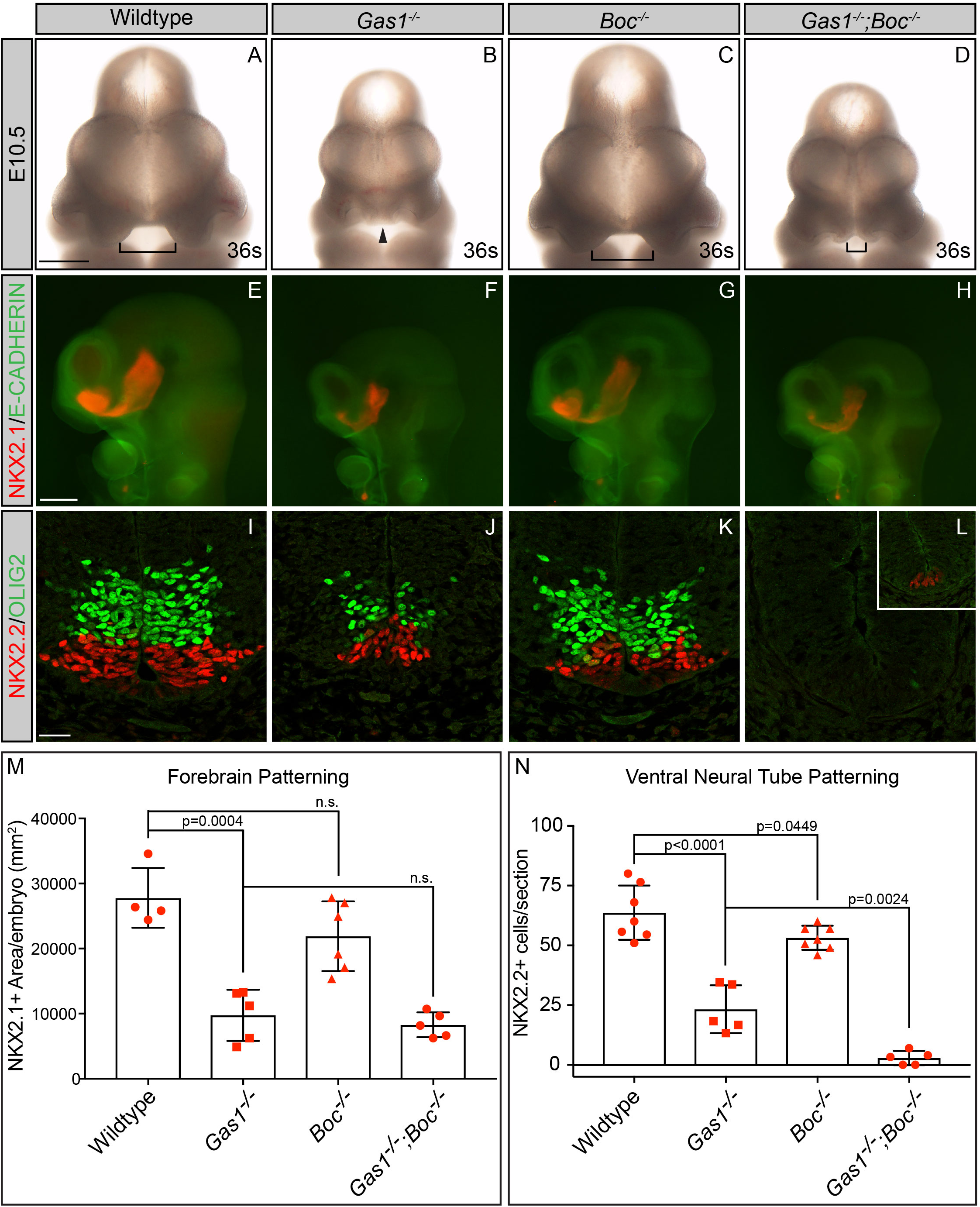
Selective contribution of *Boc* to patterning of the neural tube, but not the forebrain neuroepithelium. En face view of E10.5 embryos (A-D). Somite number (s) is indicated in the lower right corner of each panel. Brackets indicate internasal distance. Black triangles denote fusion of the MNP. E10.5 wildtype (A), *Gas1^-/-^* (B), *Boc^-/-^* (C), and *Gas1^-/-^;Boc^-/^*^-^ (D) embryos are shown. Whole-mount immunofluorescent antibody detection of E-CADHERIN (green; E-H) and NKX2.1 (red; E-H) in E10.5 wildtype (E), *Gas1^-/-^* (F), *Boc^-/-^* (G), and *Gas1^-/-^;Boc^-/^*^-^ (H) embryos. Antibody detection of OLIG2 (green; I-L) and NKX2.2 (red; I-L) in transverse sections of E10.5 forelimb level neural tubes from wildtype (I), *Gas1^-/-^* (J), *Boc^-/-^* (K), and *Gas1^-/-^;Boc^-/^*^-^ (L) embryos. Quantitation of NKX2.1 expression in wildtype (n=4), *Gas1^-/-^* (n=5), *Boc^-/-^* (n=6), and *Gas1^-/-^;Boc^-/-^* (n=5) embryos (M). Quantitation of NKX2.2+ cells (2 sections/embryo) for wildtype (n=7), *Gas1^-/-^* (n=5), *Boc^-/-^* (n=7), and *Gas1^-/-^;Boc^-/-^* (n=5) embryos (N). Data are presented as mean ± standard deviation. P-values were determined by two-tailed Student’s *t*-test. Note that NKX2.2+ cells are only present in a subset of sections from *Gas1^-/-^;Boc^-/-^* embryos (inset in L). Scale bars in (A) and (E), 500µm; (I), 25 µm.

We also analyzed HH-dependent neural patterning in the spinal cord of wildtype, *Gas1^-/-^, Boc^-/-^*, and *Gas1^-/-^;Boc^-/-^* embryos (Fig. 4 I-L,N). We examined the expression of NKX2.2 and OLIG2, two direct HH transcriptional targets that are activated in response to high and moderate levels of SHH signaling, respectively (Dessaud et al., 2008; Lei et al., 2006; Wang et al., 2011). At E10.5, *Gas1^-/-^* embryos display a significant reduction in the number of NKX2.2+ cells compared to wildtype embryos (Fig. 4J,N). Quantitation of patterning in *Boc^-/-^* embryos revealed a slight, but significant reduction in the NKX2.2 population (Fig. 4K,N). Strikingly, *Gas1^-/-^;Boc^-/-^* embryos have a very severe phenotype– OLIG2 expression is completely absent (Fig. 4L), and we observe a near complete absence of NKX2.2 expression (Fig.4 L,N). In some sections from *Gas1;Boc* mutants we could detect a few NKX2.2+ cells (Fig. 4L, inset). Overall, these data are consistent with previous studies (Allen et al., 2011), and further demonstrates that *Boc* selectively contributes to spinal cord, but not forebrain neural patterning.

Given that E10.5 *Gas1^-/-^;Boc^-/-^* mutants manifest a partial rescue of the craniofacial defects observed in *Gas1* single mutants, we investigated whether this rescue is maintained over developmental time. This question is particularly relevant since a prior analysis of *Gas1^-/-^;Boc^-/-^* embryos maintained on a mixed 129sv/C57BL/6/CD1 background demonstrated severe craniofacial defects such as clefting of the lip, palate and tongue, and disruption of the maxillary incisor (Seppala et al., 2014). To address this question, we examined craniofacial development in E18.5 wildtype and mutant embryos (Fig. 5A-D). Consistent with previous work, E18.5 *Gas1^-/-^* embryos display a range of craniofacial defects, while *Boc^-/-^* embryos appear phenotypically normal (Fig. 5A-C, Fig. S7A-B, E-F) (Allen et al., 2011; Allen et al., 2007; Seppala et al., 2007; Seppala et al., 2014; Zhang et al., 2011). *Gas1*^-/-^ and *Gas1^-/-^;Boc^-/-^* embryos share defects that include microphthalmia, midface and mandible hypoplasia, and cleft palate (Martinelli and Fan, 2007). Strikingly, and similar to what was observed during earlier developmental stages, E18.5 *Gas1^-/-^;Boc^-/-^* mutants display a less severe phenotype in specific craniofacial structures (Fig. 5D,P). Specifically, *Gas1^-/-^;Boc^-/-^* mutants display a wider maxilla and partial separation of the nasal pits; in comparison, *Gas1^-/-^* embryos have a smaller maxilla and no separation of the nasal pits (cf. black and white arrows in Fig. 5B,D). Skeletal preparations (Fig. 5E-L) confirm that *Gas1^-/-^;Boc^-/-^* mutants exhibit separation of the nasal capsule, while in *Gas1^-/-^* single mutants the nasal capsule is not separated (Fig. 5F,H). In addition to the nasal capsule, some *Gas1^-/-^;Boc^-/-^* embryos exhibit widening of the premaxilla, although in others it is hypoplastic (see red arrow in Fig. 5H and inset in Fig. S7H). These data suggest that the amelioration of the craniofacial defects observed at E10.5 in *Gas1;Boc* mutant embryos persists over developmental time.

**Figure 5.**
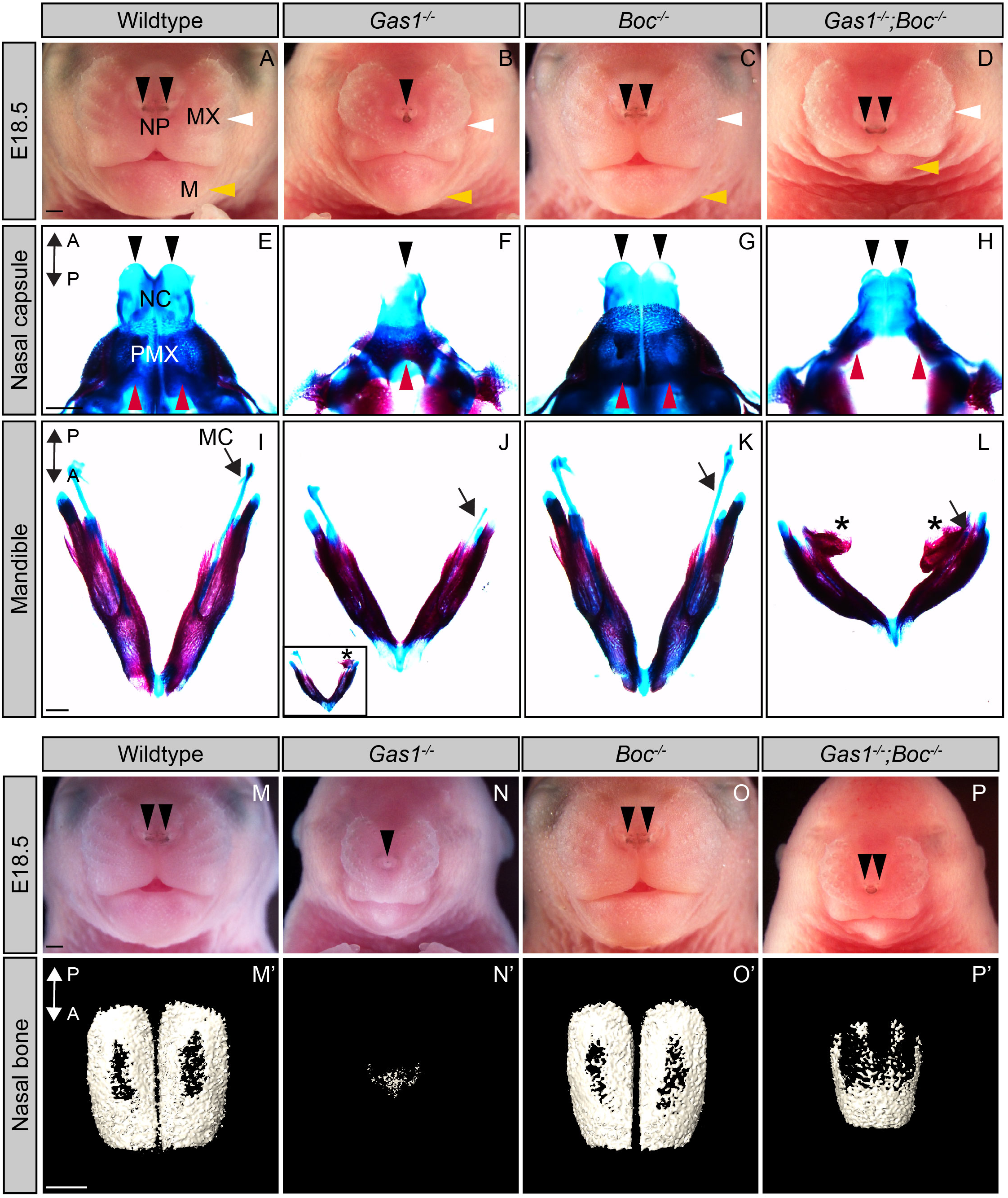
Partial rescue of HPE phenotypes persists through E18.5 in *Gas1;Boc* mutant embryos. En face view of E18.5 wildtype (A,M), *Gas1^-/-^* (B,N), *Boc^-/-^* (C,O), and *Gas1^-/-^;Boc^-/-^* (D,P) embryos. Black arrowheads denote the nasal pits (NP), white arrowheads mark the maxilla (MX), and yellow arrowheads identify the mandible (M). E18.5 craniofacial structures stained with alcian blue and alizarin red to visualize cartilage and bone, respectively (E-L). Dorsal views of the nasal capsule (NC) and premaxilla (PMX) of E18.5 wildtype (E), *Gas1^-/-^* (F), *Boc^-/-^* (G), and *Gas1^-/-^;Boc^-/-^* (H) are shown. Black arrowheads indicate the nasal capsule and red arrowheads mark the premaxilla. Dorsal views of the mandible of E18.5, wildtype (I), *Gas1^-/-^* (J), *Boc^-/-^* (K), and *Gas1^-/-^;Boc^-/-^* (L) are shown. Asterisks identify ectopic bone duplications in the posterior part of the mandible and black arrows denote Meckel’s cartilage (MC). Inset in J, shows ectopic bone in a *Gas1^-/-^* mutant embryo. Three dimensional reconstructions of microCT images of isolated nasal bones from E18.5 wildtype (M’), *Gas1^-/-^* (N’), *Boc^-/-^* (O’), and *Gas1^-/-^;Boc^-/-^* (P’) embryos. A←→P specifies the anterior to posterior axis in (E-H, I-L, M’-P’). Scale bars (A, E, I, M, M’), 500µm.

In contrast to the nasal capsule and premaxilla, *Gas1^-/-^;Boc^-/-^* embryos exhibit a shortened mandible and truncated meckel’s cartilage compared to *Gas1^-/-^* embryos (Fig. 5J,L). The mandible of *Gas1^-/-^;Boc^-/-^* mutants also exhibit ectopic bone duplications on the posterior inferior side of the mandible (Fig. 5L). Occasionally, *Gas1^-/-^* mutants with severe HPE phenotypes display a similar phenotype (Fig. 5J inset). Bone duplications have been associated with loss of HH signaling in the mandibular neural crest-derived mesenchyme (Jeong et al., 2004; Xu et al., 2019). *Gas1^-/-^;Boc^-/-^* mutants also display severe defects in the maxilla, palatine bone and the occipital bone (Fig. S7H). We also evaluated SHH-dependent digit specification in these embryos (Fig. S7E’-H’). Consistent with previous work (Allen et al., 2011), combined loss of *Gas1* and *Boc* results in severe digit specification defects (Fig. S7H’). These results suggest opposing and tissue-specific contributions of *Boc* to HH-dependent craniofacial development.

To further investigate these phenotypes, we analyzed three (3D) dimensional reconstructions from micro-computed tomography (µCT) images (Fig. 5M’-P’, Fig. S7A’-D’). Specifically, we focused on the nasal bone, where we observed the partial rescue in *Gas1;Boc* mutants. The 3D reconstructions indicated that the nasal bone in *Gas1^-/-^;Boc^-/-^* mutants is partially restored compared to *Gas1^-/-^* mutants where this bone is smaller and fragmented (Fig. 5N’, P’). As we observed at E10.5 (Fig. 2), there is a spectrum of HPE phenotypes in *Gas1* mutants (Fig. S7A-D); however, we consistently observe that the nasal bone of *Gas1*;*Boc* mutants is not as severely affected as the *Gas1* single mutants (Fig. S7A’-D’). These data confirm that *Gas1^-/-^;Boc^-/-^* embryos display a less severe phenotype in the nasal bone and in the nasal capsule than *Gas1^-/-^* embryos.

To investigate the mechanisms that could explain the partial rescue observed in *Gas1^-/-^;Boc^-/-^* embryos, we analyzed tissue-specific proliferation in the telencephalon of E10.5 wildtype and mutant embryos. Specifically, we performed immunofluorescent detection of Phospho-Histone H3 (PH3) in slides and co-stained with antibodies directed against E-CADHERIN (E-CAD) and PDGFRα to discriminate between the surface ectoderm, forebrain neuroepithelium, and craniofacial mesenchyme (Fig. 6A-E). Coronal sections of E10.5 *Gas1^-/-^* mutant embryos display reduced numbers of PH3+ cells across the surface ectoderm, forebrain neuroepithelium, and craniofacial mesenchyme (Fig. 6B, F-H). Similarly, *Cdon^-/-^* embryos exhibit a significant decrease in proliferation both the surface ectoderm and craniofacial mesenchyme (Fig. 6C, F-H). In contrast, *Boc^-/-^* embryos do not display any apparent changes in the proliferation in the surface ectoderm or in the neuroepithelium (Fig. 6D, F,G). Further, *Boc^-/-^* embryos display a significant increase in mesenchymal proliferation compared to wildtype embryos (Fig. 6H). These results suggest that *Boc* negatively regulates proliferation specifically in craniofacial mesenchyme.

**Figure 6.**
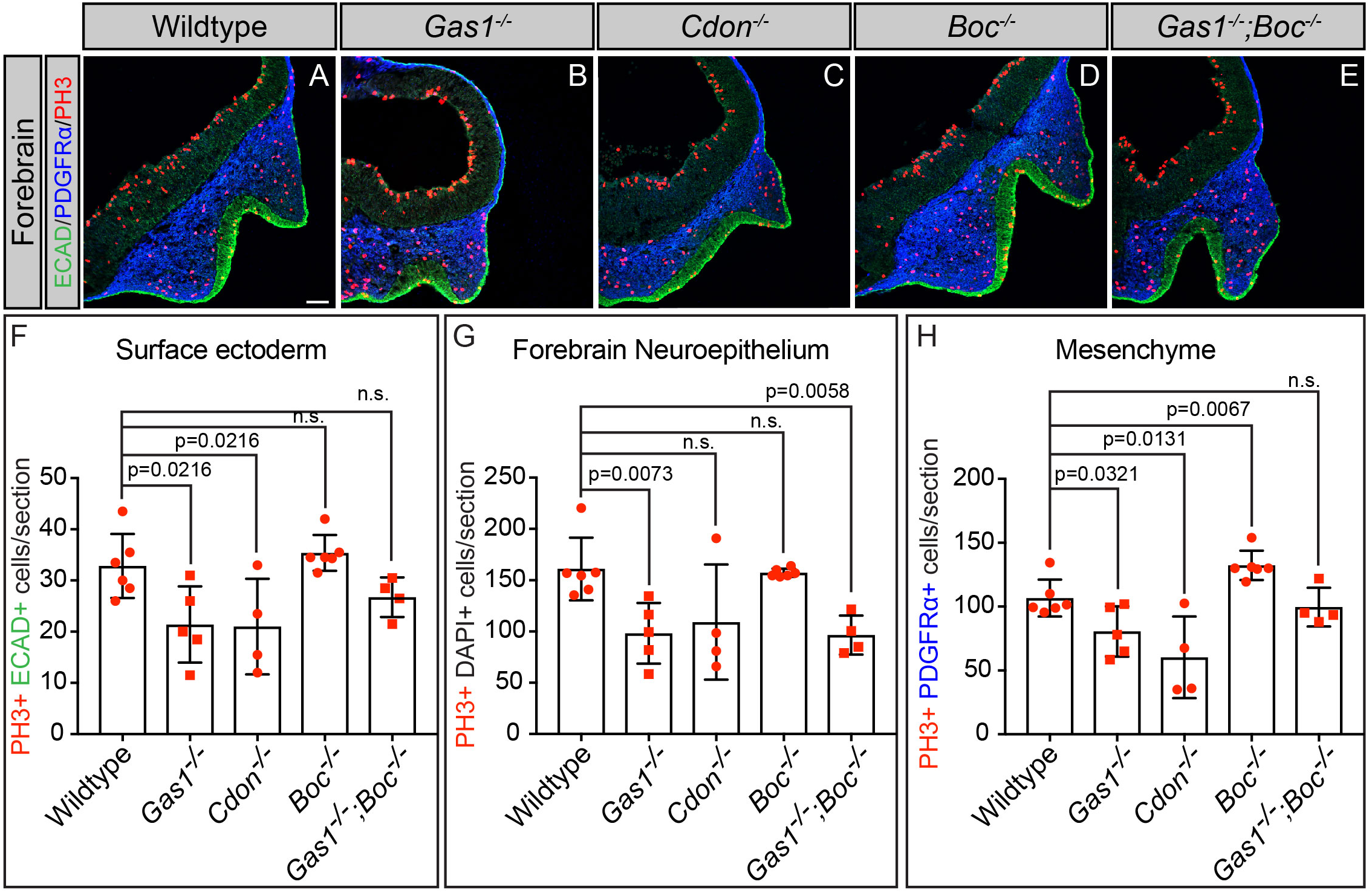
*Boc* selectively inhibits mesenchymal proliferation during craniofacial development. Immunofluorescent analysis of proliferation in E10.5 telencephalon coronal sections from wildtype (A), *Gas1^-/-^* (B), *Cdon^-/-^* (C), *Boc^-/-^* (D), and *Gas1^-/-^;Boc^-/-^* (E) embryos. Antibody detection of E-CADHERIN (ECAD, green), PDGFRα (blue) and phospho-histone H3 (PH3, red). Quantitation of PH3+ cells (2 sections/embryo) in the surface ectoderm (F), forebrain neuroepithelium (G), and craniofacial mesenchyme (H), of E10.5 wildtype (n=6), *Gas1^-/-^* (n=5), *Boc^-/-^* (n=6), and *Gas1^-/-^;Boc^-/-^* (n=4). Data are presented as mean ± standard deviation. P-values were determined by two-tailed Student’s *t*-test. Note that *Boc^-/-^* embryos display increased proliferation in the craniofacial mesenchyme (H). Scale bar (A), 50 µm.

We also investigated tissue-specific proliferation in *Gas1^-/-^;Boc^-/-^* mutant embryos. Notably, the levels of proliferation in the surface ectoderm and the mesenchyme are not significantly different when compared to wildtype embryos (Fig. 6F,H) In contrast, proliferation is significantly decreased in the forebrain neuroepithelium of *Gas1;Boc* mutants (Fig. 6G). Surprisingly, this effect on proliferation appears to be quite selective, as there are no significant changes in proliferation in *Boc* mutants in either the neural tube or the forelimb mesenchyme (Fig. S8). Overall, these data demonstrate that *Boc* functions in a non-redundant manner to restrict proliferation in the craniofacial mesenchyme, while acting in concert with *Gas1* and *Cdon* to promote proliferation in the forebrain neuroepithelium.

## Discussion

Here we investigated the individual and combined contributions of the HH co-receptors *Gas1*, *Cdon* and *Boc* during HH-dependent craniofacial development. We found that *Boc* displays a significantly broader expression pattern than *Gas1* and *Cdon* in multiple craniofacial structures. Surprisingly, and distinct from *Gas1* and *Cdon*, loss of *Boc* alone does not result in any detectable reduction of HH pathway activity in developing craniofacial structures. Instead, we find that genetic deletion of *Boc* results in facial widening that is consistent with increased HH pathway activity (Brugmann et al., 2010; Hu and Helms, 1999). Further, analysis of *Gas1;Boc* double mutants revealed an amelioration of the craniofacial phenotype observed in *Gas1* single mutants, corresponding with increased HH pathway activity, and consistent with the notion that loss of *Boc* can counterintuitively drive increased HH signaling. Notably, this improvement is restricted to a subset of craniofacial structures, but persists throughout embryonic development. Mechanistic analyses suggest that *Boc* achieves these tissue-specific effects through the selective restriction of proliferation in the neural crest-derived mesenchyme. Taken together, these data demonstrate that *Boc* regulates HH signaling in a tissue-specific manner, and suggests that, in certain tissues, BOC works in opposition to other HH co-receptors to restrain HH pathway function.

### Genetic background-dependent phenotypic differences in HH co-receptor mutants

Understanding the molecular mechanisms that underlie HPE is confounded by the significant phenotypic variability observed in this disease, and the complex genetics that contribute to proper craniofacial development. Our data indicate that, even when maintained on a congenic C57BL/6J background, *Gas1* and *Cdon* mutants display a range of HPE phenotypes. These phenotypes vary from microforms of HPE to semilobar HPE, and their severity correlates with HH pathway activity as assessed by *Gli1* expression. The variability in the HPE phenotypes of our mutants could be explained due to multiple genetic and non-genetic risk factors (Hong and Krauss, 2018). In particular, the variable severity across the phenotypes in our mutants could arise from stochastic changes in the establishment or response to the SHH morphogen gradient in the neuroepithelium, neural crest-derived mesenchyme, and/or surface ectoderm. In early craniofacial structures *Shh* is expressed sequentially, initiating in the prechordal plate, followed by the diencephalon and telencephalon, subsequently in the surface ectoderm of the frontonasal prominence, and finally in the pharyngeal endoderm of the first branchial arch (Aoto et al., 2009; Cordero et al., 2004; Marcucio et al., 2005; Rubenstein and Beachy, 1998; Xavier et al., 2016a). This complex developmental expression sequence of *Shh*, which is required to properly pattern the craniofacial structures (Krauss, 2007), combined with the differential expression of multiple HH receptors could generate an inherent variability that affects the severity of the HPE phenotypes.

The lack of craniofacial defects in *Boc* mutants maintained on different genetic mixed backgrounds (Okada et al., 2006; Seppala et al., 2014; Zhang et al., 2011) suggested a minor, redundant role for *Boc* in HH-dependent craniofacial development. This notion of *Boc* as a silent HPE modifier gene is supported by studies where *Boc* deletion in a *Gas1* or *Cdon* null background enhances HPE severity and decreases the levels of HH pathway targets (Seppala et al., 2014; Zhang et al., 2011). However, our data indicate that *Boc* mutants on a C57BL/6J background exhibit internasal distance widening in E10.5 embryos. These data suggest an antagonistic role for *Boc* in HH signaling and comports with a previous description of *Boc* as a potential HH pathway antagonist in the zebrafish lower jaw (Bergeron et al., 2011). While we do not observe any mandible phenotypes in *Boc^-/-^* embryos, species-specific differences in craniofacial development between mouse and fish likely limit our ability to draw a direct connection. Alternatively, our analysis of *Boc* in the developing mandible may not be comprehensive enough to reveal this function. Regardless, our data reveal a novel, antagonistic role for *Boc* during aspects of craniofacial development, and raises the question of whether BOC may work in concert with other known redundant HH pathway antagonists, including PTCH1, PTCH2 and HHIP1, to maintain the balance between HH pathway activation and inhibition (Holtz et al., 2013) in craniofacial structures. Additionally, our data suggest that HH co-receptors can function to alternately promote or antagonize HH signaling depending on the context. In support of this notion, *Gas1* (Cobourne et al., 2004; Lee et al., 2001; Ohazama et al., 2009) and *Cdon* (Cardozo et al., 2014) can negatively regulate HH pathway function in different tissues.

*Boc* deletion partially rescues the HPE phenotypes of *Gas1* single mutants. Specifically, *Gas1;Boc* double mutants display increased MNP separation at E10.5, and restoration of the nasal capsule and nasal bone at E18.5. Importantly, these phenotypes correlate with increased *Gli1* levels, suggesting that *Boc* selectively antagonizes HH signaling during craniofacial development. These data partially contrast with previous work (Seppala et al., 2014), in which *Gas1;Boc* mutants on a 129Sv-C57BL/6/CD1 genetic background display more severe phenotypes than those observed in *Gas1* mutants (Seppala et al., 2014). Although *Gas1;Boc* mutants on a C57BL/6J background display severe defects in the majority of the bones of the skull and cleft palate as previously reported (Seppala et al., 2014), we never observe clefting of the lip in these mutants. Given that the lip is formed by the fusion of the MXP and MNP (Jiang et al., 2006), this result is consistent with the partial rescue mediated by *Boc* deletion in the nasal bone and nasal capsule.

### Tissue-specific functions of BOC in HH signal transduction

Analysis of HH transcriptional targets revealed that *Boc* deletion results in differential changes in HH-dependent gene expression in a tissue-specific fashion (Fig. 7A). Specifically, our data suggest that BOC promotes the expression of the direct HH transcriptional target, NKX2.2, in the spinal cord neuroepithelium, but does not contribute to expression of NKX2.1 in the telencephalon neuroepithelium. These data suggest that BOC differentially regulates HH-dependent neural patterning at distinct axial levels. Further, BOC promotes *Gli1* expression in the limb bud mesenchyme, but antagonizes *Gli1* expression in the forebrain mesenchyme. Notably, *Boc* appears to selectively impact HH-dependent patterning, but not proliferation in the developing limb bud; conversely, *Boc* selectively inhibits proliferation in the neural crest-derived mesenchyme of the craniofacial structures (Fig 7A). This is consistent with previous work by (Xavier et al., 2016b) suggesting that *Boc* contributes to mesenchymal proliferation in the palatal shelf. Taken together, these data argue that BOC regulates patterning and proliferation in a tissue-specific manner and raises the possibility that BOC performs multiple, and in some cases, opposing roles in HH signal transduction.

**Figure 7.**
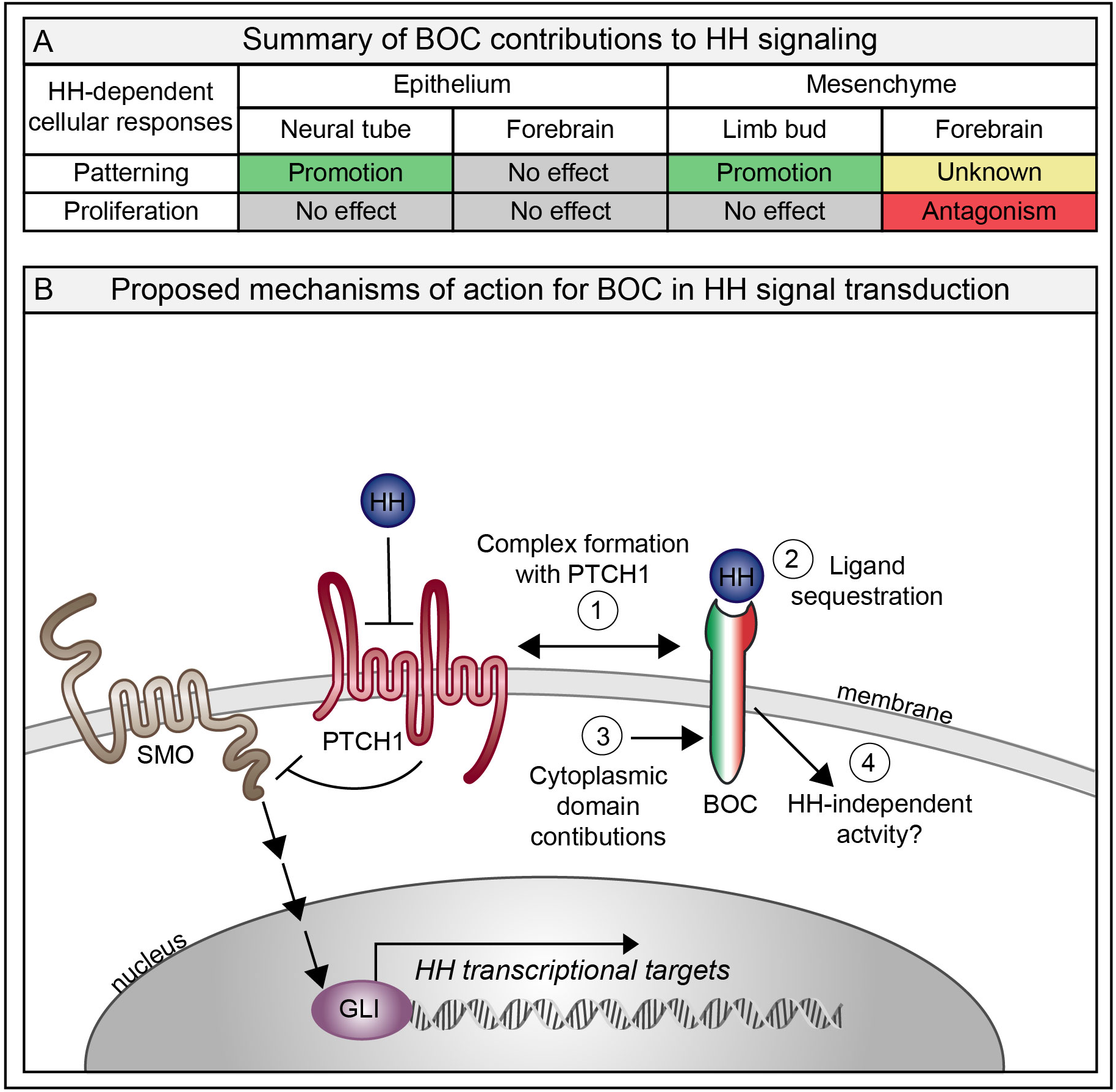
BOC is a multi-functional regulator of HH signaling. Summary of BOC contributions to HH signaling (A). Green indicates promotion of HH signaling, red denotes HH pathway antagonism, gray suggests no effect, and yellow is unknown. Proposed mechanisms of action for BOC in HH signal transduction (B). 1. Complex formation with PTCH1.The interaction of PTCH1 and BOC that allows the formation of a receptor complex that alternately activates or inhibits HH pathway activity. 2. Ligand sequestration. BOC binds HH ligands through its extracellular domain and could antagonize HH signaling by sequestering SHH in areas of low SHH concentration. 3. Cytoplasmic domain contributions. The unique cytoplasmic domain of BOC could regulate additional downstream signaling cascades that enable its tissue-specific functions. 4. HH-independent activity. BOC could mediate yet to be identified HH-independent functions that either augment or counter the HH response.

### Boc as a multi-functional regulator of HH signaling

Based on our data, and the work of others, we propose a model whereby BOC acts as a multi-functional receptor to contribute to vertebrate embryogenesis (Fig. 7B). Specifically, we propose that BOC can act to: 1) promote HH signaling through interactions with HH ligands and the canonical receptor PTCH1; 2) antagonize HH signaling, either through ligand sequestration, or perhaps through the formation of an inhibitory complex with PTCH1; 3) contribute to HH-dependent signaling via its unique cytoplasmic domain; 4) function independently of the HH pathway.

BOC physically interacts with PTCH1 in a SHH-independent manner (Izzi et al., 2011). In craniofacial structures PTCH1 and BOC are co-expressed in a subset of cells in the MNP (Seppala et al., 2014). The differential interaction of these proteins could allow the formation of a receptor complex that alternately activates or inhibits HH pathway activity. Alternatively, BOC binding to HH ligands via its extracellular domain (Beachy et al., 2010; McLellan et al., 2008; Yao et al., 2006) raises the possibility that BOC can sequester SHH ligand in areas of low SHH concentration, and subsequently antagonize HH signaling. Consistent with this notion, *Boc* expression in HH-responsive tissues generally extends closer to the source of SHH ligand than either *Gas*1 or *Cdon*.

BOC displays a unique cytoplasmic domain that does not resemble any other protein or motif (Kang et al., 2002). Recently work suggests that the BOC cytoplasmic domain binds to the non-receptor tyrosine kinase ABL (Vuong et al., 2017) and to the adaptor protein ELMO1 (Makihara et al., 2018). Thus, this domain could be critical to mediate tissue-specific, HH-dependent signals, or to perform HH-independent functions through the activation of downstream signaling cascades. It will be interesting to investigate the contribution of the BOC cytoplasmic domain to its tissue-specific functions during craniofacial development. Overall, this work identifies multiple and distinct roles for BOC in HH-dependent craniofacial development.

## Materials and methods

### Reagents

For reagents and primary antibodies see supplemental table 1, and supplemental table 2, respectively in the supplementary information.

### Animal Models

*Gas1^lacZ^* (Martinelli and Fan, 2007), *Cdon^lacZ-2^* (Cole and Krauss, 2003), and *Boc^AP^*(Zhang et al., 2011) mice have been all described previously. *Gas1, Cdon,* and *Boc* mutants were backcrossed for at least ten generations to create lines on a congenic C57BL/6J background. *Cdon^lacZ-1^* mice (Cole and Krauss, 2003) were maintained on a mixed 129/Sv/C57BL/6 background for expression analysis. For embryonic dissections, noon of the day on which a vaginal plug was detected was considered as E0.5. For precise staging, somites were counted during the dissection. Embryos with 34-38 somites were considered E10.5 embryos. Fertilized eggs were obtained from the Poultry Teaching & Research Center at Michigan State University. To obtain Hamburger-Hamilton (HH) stage 11 chicken embryos, the fertilized eggs were incubated 39-40 hours at 37°C in a GQF 1550 hatcher incubator with normal humidity settings (45%-55%). All animal procedures were reviewed and approved by the Institutional Animal Care and Use Committee (IACUC) at the University of Michigan.

### X-gal staining

Embryos were dissected in 1X PBS, pH 7.4, and fixed (1% formaldehyde, 0.2% glutaraldehyde, 2mM MgCl_2_, 5mM EGTA, 0.02% NP-40) on ice for 10-60 minutes depending on the embryonic stage. Subsequently, the embryos were washed 3 x 5 minutes with 1X PBS, pH 7.4 + 0.02% NP-40 for permeabilization. B-Galactosidase activity was detected with X-Gal staining solution (5mM K_3_Fe(CN)_6_, 5mM K_4_Fe(CN)_6_, 2mM MgCl_2_, 0.01% Na deoxycholate, 0.02% NP-40, 1mg/mL X-gal). The signal was developed from 25 minutes to 24 hours at 37° C depending on the *lacZ* allele. After staining, the embryos were washed 3 x 5 minutes with 1X PBS, pH 7.4 at 4°C, and post-fixed in 4% paraformaldehyde for 20 minutes at room temperature, followed by 3 x 5 minute washes in 1X PBS, pH 7.4. Finally, embryos were stored and photographed in 1X PBS, pH 7.4 + 50% glycerol. X-gal staining of sections (20µm) was performed as described above for whole mount embryos. After staining, sections were washed 3 x 5 minutes with 1X PBS, pH 7.4, counterstained with nuclear fast red for 5 minutes and dehydrated in an ethanol series (70% ethanol, 95% ethanol, 100% ethanol and 100% Xylenes) followed by application of coverslips with permount mounting media.

### Alkaline Phosphatase Staining

Embryos were dissected on 1X PBS, pH 7.4, and fixed (1% formaldehyde, 0.2% glutaraldehyde, 2mM MgCl_2_, 5mM EGTA, 0.02% NP-40) on ice for 10-60 minutes depending on the embryonic stage on ice. Subsequently, the embryos were washed 3 x 5 minutes with 1X PBS, pH 7.4. To deactivate endogenous alkaline phosphatases, embryos were incubated in 1X PBS, pH 7.4 at 70°C for 30 minutes. Then the embryos were rinsed with 1X PBS, pH 7.4 and washed for 10 minutes in alkaline phosphatase buffer (100mM NaCl, 100mM Tris-HCl pH9.5, 50mM MgCl_2_, 1% Tween-20) at room temperature. Embryos were stained with BM purple from 2 to 3 hours at 37°C depending on the embryonic stage. After staining, the embryos were washed 3 x 5 minutes with 1X PBS, pH 7.4 at 4°C, and post-fixed in 4% paraformaldehyde for 20 minutes at room temperature, followed by 3 x 5 minute washes with 1X PBS, pH 7.4. Finally, embryos were stored and photographed in 1X PBS, pH 7.4 + 50% glycerol. Alkaline phosphatase staining of sections (20µm) was performed as described above for whole mount embryos. After staining, sections were washed 3 x 5 minutes with 1X PBS, pH7.4, counterstained with nuclear fast red for 5 minutes and dehydrated in an ethanol series (70% ethanol, 95% ethanol, 100% ethanol and 100% xylenes for five minutes each) followed by application of coverslips with permount mounting media.

### Whole-Mount Digoxigenin *in situ* Hybridization

Whole-mount digoxigenin *in situ* hybridization was performed as previously described in (Allen et al., 2011; Wilkinson, 1992). In brief, embryos were dissected in 1X PBS, pH 7.4 and fixed in 4% paraformaldehyde overnight on a rocking platform. After fixation, embryos were dehydrated in a methanol/PBST (1X PBS, pH 7.4 + 0.1 % Tween) series (25% methanol, 50 %methanol, 75% methanol) and stored in 100% methanol at −20°C until the experiment was performed for up to 6 months. Embryos were digested with 10µg/mL proteinase K at RT for 2 minutes. Hybridization was performed with the indicated digoxigenin probe with a concentration of 1ng/µL for 16-19 hours at 70°C. The embryos were incubated in alkaline phosphatase-conjugated anti-DIG antibody at a dilution of 1:4,000. AP-anti-DIG was detected with BM purple, and signal was developed for 3.5 hours at room temperature. Embryos were cleared in 50% glycerol in 1XPBST and were photographed using a Nikon SMZ1500 microscope.

### Immunofluorescence

Section immunofluorescence was performed as in (Allen et al., 2011). Embryos were dissected in 1X PBS, pH 7.4 and fixed for 1 hour in 4% paraformaldehyde on ice, followed by 3 x 5 minutes washes with 1X PBS, pH 7.4 and cryoprotected for 24-48 hours in 1X PBS + 30% sucrose. Embryos were embedded in OCT compound and sectioned on a Leica cryostat (12 µm thick forebrain and forelimb neural tube sections). Sections were blocked in blocking buffer (3% bovine serum albumin, 1% heat-inactivated sheep serum, 0.1% TritonX-100 in 1X PBS, pH 7.4) for 1 hour. Primary antibodies were diluted in blocking buffer incubated overnight at 4 °C in a humidified chamber. A list of all the primary antibodies used in this study is provided in supplementary table 2. Secondary antibodies were diluted in blocking solution and incubated for 1 hour at room temperature, followed by 3 x 5 minute washes with 1X PBS, pH 7.4. All Alexa Fluor Dyes secondary antibodies were used at a 1:500 dilution. Nuclei were labeled with DAPI for 10 minutes at room temperature and slides were mounted with coverslips using Immu-mount aqueous mounting medium. Sections were visualized on a Leica upright SP5X confocal microscope.

### Whole-Mount Immunofluorescence

Embryos were dissected in 1X PBS, pH 7.4, fixed with 4% paraformaldehyde for 2 hours at 4°C, and washed 2 x 10 minutes washes with PBTX (1X PBS + 0.1% Triton X-100). Subsequently, embryos were blocked for 1 hour in PBTX + 10% goat serum. Primary antibodies were diluted in PBTX + 10% goat serum and incubated overnight at 4 °C on a rocking platform. A list of all the primary antibodies used in this study is provided in the supplementary table 2. The next day the embryos were rinsed 2 x 5 minutes with PBTX, followed by 3 x 1 hour washes with PBTX on a rocking platform at 4°C. After the washes, embryos were incubated overnight with secondary antibodies diluted in PBTX+ 10% serum. All Alexa Fluor Dyes secondary antibodies were used at a 1:500 dilution. Next, embryos were washed as described for the primary antibody above, and cleared with *Clear^T2^* (25% Formamide/10%PEG for one hour; 50% Formamide/20%PEG for 72 hours) (Kuwajima et al., 2013). Finally, embryos were visualized on a Nikon SMZ1500 microscope. With the *Clear^T2^* reagent we did not observed any tissue expansion. (Protocol courtesy of Jean-Denis Benazet, UCSF)

### Micro Computed Tomography (Micro CT)

E18.5 embryos were skinned and eviscerated. Subsequently, embryos were fixed overnight in 100% ethanol, and maintained in 70% ethanol until ready to scan. The scans were performed using embryos covered with a 1X PBS, pH 7.4-soaked kim wipe and scanned over the entire length of the skull using the µCT100 system (Scanco Medical, Bassersdorf, Switzerland). Scan settings were as follows: 12 µm voxel size, 55 kVp, 109 µA, 0.5 mm AL filter, and 500 ms integration time. Micro CT scans were analyzed with the Amira software (Thermo Fisher Scientific). The Micro CT scans were uploaded as DICOM files into the software and the three-dimensional reconstructions were generated using the isosurface feature. The individual bones were manually segmented using the extract surface and buffer tools of Amira (Ho et al., 2015). Finally, the individual bones were color coded.

### Skeletal Preparation

Skeletons were prepared as previously described before in (Allen et al., 2011). E18.5 embryos were skinned and eviscerated. Subsequently, embryos were fixed in 100% ethanol, followed by 100% acetone for 24 hours respectively at room temperature. Cartilage and bone were stained with alcian blue/alizarin red staining solution (5% alcian blue, 5% alizarin red, 5% glacial acetic acid and 70% ethanol) for 4 days at room temperature. The remaining tissue was digested with several washes of 1% potassium hydroxide. The skeletons were cleared by 24 hour washes of a gradient of glycerol (20%, 50%, and 80%) in 1% potassium hydroxide, and photographed in 80% glycerol.

### *In ovo* chicken electroporations

Chicken electroporations were performed as previously described in (Allen et al., 2011; Tenzen et al., 2006). The indicated construct (pCIG plasmid −1 µg/µl in 1X PBS, pH7.4, with 50ng/µl fast green) was injected into the forebrain cavity of HH stage 11 chicken embryos. L-shaped electrodes were made with platinum wire, 8mm long (3mm were bent to form the L shape) and spaced 6mm apart. Electrodes (L-shaped part) were placed in front of the forebrain of the embryo (pulsed five times at 25 V for 50 ms with a BTX electroporator). The electroporated embryos were screened for GFP expression after 48 hours at HH stage 21-22 and processed for immunofluorescence.

### Quantitation and statistical analysis

All the data are represented as mean ± standard deviation. All statistical analyses were performed using GraphPad statistic calculator (GraphPad Software, La Jolla California USA, www.graphpad.com). Statistical significance was determined using two-tailed Student’s *t-*test. Significance was defined according GraphPad Prism style: non-significant (p>0.05), * (p≤0.05), **(p≤0.01), ***(p≤0.001) and ****(p≤0.0001). For all the experimental analyses a minimum of 3 embryos of each genotype were examined, each n represents an embryo. All the statistical details (statistical tests used, statistical significance and exact value of each n) for each experiment are specified in the figure legends.

### Telencephalic division and medial nasal process classification

Frontal pictures of E10.5 mouse embryos were photographed with a Nikon SMZ1500 microscope. Blind classification of the telencephalic division and media nasal process separation, was performed by a blinded evaluator according the categories showed in (Fig S2A-F).

### Internasal distance and crown-rump length quantitation

Pictures of the nasal processes and whole E10.5 embryos were taken in 1X PBS, pH7.4 with a Nikon SMZ1500 microscope. Internasal distance was defined as the distance between the edges of the medial nasal process. Crown rump length was defined as top of the crown of the midbrain, bisecting the forelimb bud to the curvature at the bottom c-shaped part of the embryo. Blind quantitation of the intranasal distance and crown-rump length was performed manually by a single evaluator using the scale bar tool of the NIS-Elements software (Nikon) annotations and measurements feature.

### Immunofluorescence quantitation

To quantify immunofluorescence images, we examined a minimum of 3 embryos per genotype and 2 sections from each embryo.

NKX2.1 quantitation: Side view pictures of whole mount immunofluorescent wildtype and mutant embryos were taken in *Clear^T2^* with a Nikon SMZ1500 microscope. The NKX2.1 area of expression was quantified using the area measure plugin of ImageJ (Schneider et al., 2012). Each image was thresholded automatically by ImageJ before the area of expression was quantified.

NKX2.2 quantitation: Pictures of transverse sections of wildtype and mutant neural tubes stained with antibodies directed against NKX2.2 were merged with their respective DAPI images. NKX2.2 positive cells were quantified with the point tool of ImageJ (Schneider et al., 2012).

Phospho-histone H3 quantitation: All phospho-histone H3 quantitation was performed with the point tool feature of ImageJ (Schneider et al., 2012). In the forebrain, the phospho-histone H3 positive cells were quantified in different tissue compartments. The phospho-histone H3 images were merged with markers specific to each tissue: E-CADHERIN (surface ectoderm), and PDGFRα (mesenchyme). The neuroepithelium was identified morphologically. The dorsal telencephalic midline was excluded from this analysis. For the neural tube quantitation, the phospho-histone H3 cells were quantified in the ventral limit of expression of the NKX6.1 neural progenitors. Finally, in the forelimb bud, the phospho-histone H3 positive cells were quantified specifically in a selected area of equal size in wildtype and mutant embryos.

## Acknowledgements

We thank all current and past members of the Allen lab for valuable feedback and suggestions throughout the course of this study. In particular, we thank Nicole Franks and Savannah Struble for significant technical assistance. We also thank Michelle Lynch (University of Michigan) for assistance with scanning MicroCT samples and Thach-Vu Ho (University of Southern California) for assistance with generating the MicroCT 3D reconstructions. We also gratefully acknowledge the Department of Cell and Developmental Biology, including the Engel, Spence, and O’Shea laboratories at the University of Michigan for providing access to research equipment. The NKX2.2 and NKX6.1 antibodies were obtained from the Developmental Studies Hybridoma Bank, created by the NICHD of the NIH and maintained at The University of Iowa, Department of Biology, Iowa City, IA 52242. Finally, we acknowledge the Biomedical Research Core Facilities Microscopy Core for providing access to confocal microscopy equipment, which is supported by the Rogel Cancer Center.

## Data Availability

This study did not generate/analyze any datasets.

## Supplemental figure legends

**Supplemental Figure 1.**
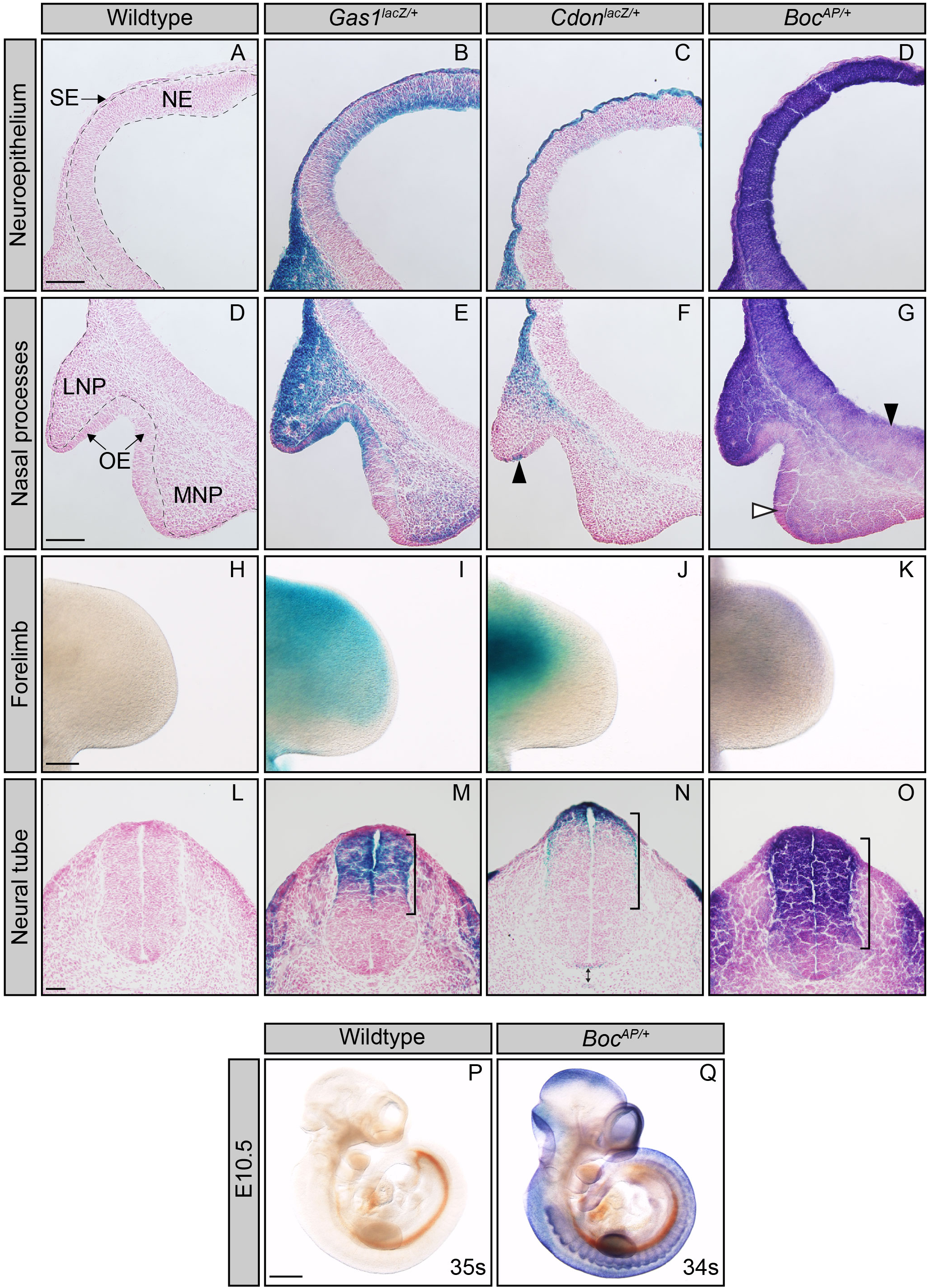
G*a*s1*, Cdon* and *Boc* are differentially expressed across multiple HH-responsive tissues. Analysis of HH co-receptor expression using *lacZ* (*Gas1*, *Cdon*) and *hPLAP* (*Boc*) reporter alleles in HH-responsive tissues (A-O). High magnification pictures of coronal sections of E10.5 telencephala (A-G; cf. Fig.1Q-T), from wildtype (A, D), *Gas1^lacZ/+^* (B, E), *Cdon^lacZ/+^* (C, F), and *Boc^AP/+^* (D, G) embryos is shown. E10.5 forebrain neuroepithelia (A-D) and nasal processes (D-G). Arrowhead in (F) denotes a subset of cells expressing *Cdon* in the olfactory epithelium. Black arrowhead in (G) identifies the extended ventral expression of *Boc* closer to the source of *Shh* expression. White arrowhead in (G) denotes *Boc* expression in the olfactory epithelium. Whole mount X-Gal and Alkaline Phosphatase staining of E10.5 forelimb buds (H-K), wildtype (H), *Gas1^lacZ/+^* (I), *Cdon^lacZ/+^* (J), and *Boc^AP/+^* (K). Tranverse sections of E10.5 neural tubes (L-O), wildtype (L), *Gas1^lacZ/+^* (M), *Cdon^lacZ/+^* (N), and *Boc^AP/+^* (O). Black brackets denote the expression domain of the HH co-receptors in the neural tube. Double-headed arrow in (N) indicates *Cdon* expression in the floor plate and notochord. Heat inactivation of endogenous alkaline phosphatase at E10.5 in wildtype (P) and *Boc^AP/+^* (Q) animals demonstrates the specificity of alkaline phosphatase staining. Somite number (s) is indicated in the lower right corner (P-Q). Scale bars, (A-G) 100µm, (H-K) 200µm, (L-O) 50µm, (P-Q) 500µm. Abbreviations: surface ectoderm (SE), neuroepithelium (NE), lateral nasal process (LNP), medial nasal process (MNP), olfactory epithelium (OE).

**Supplemental Figure 2.**
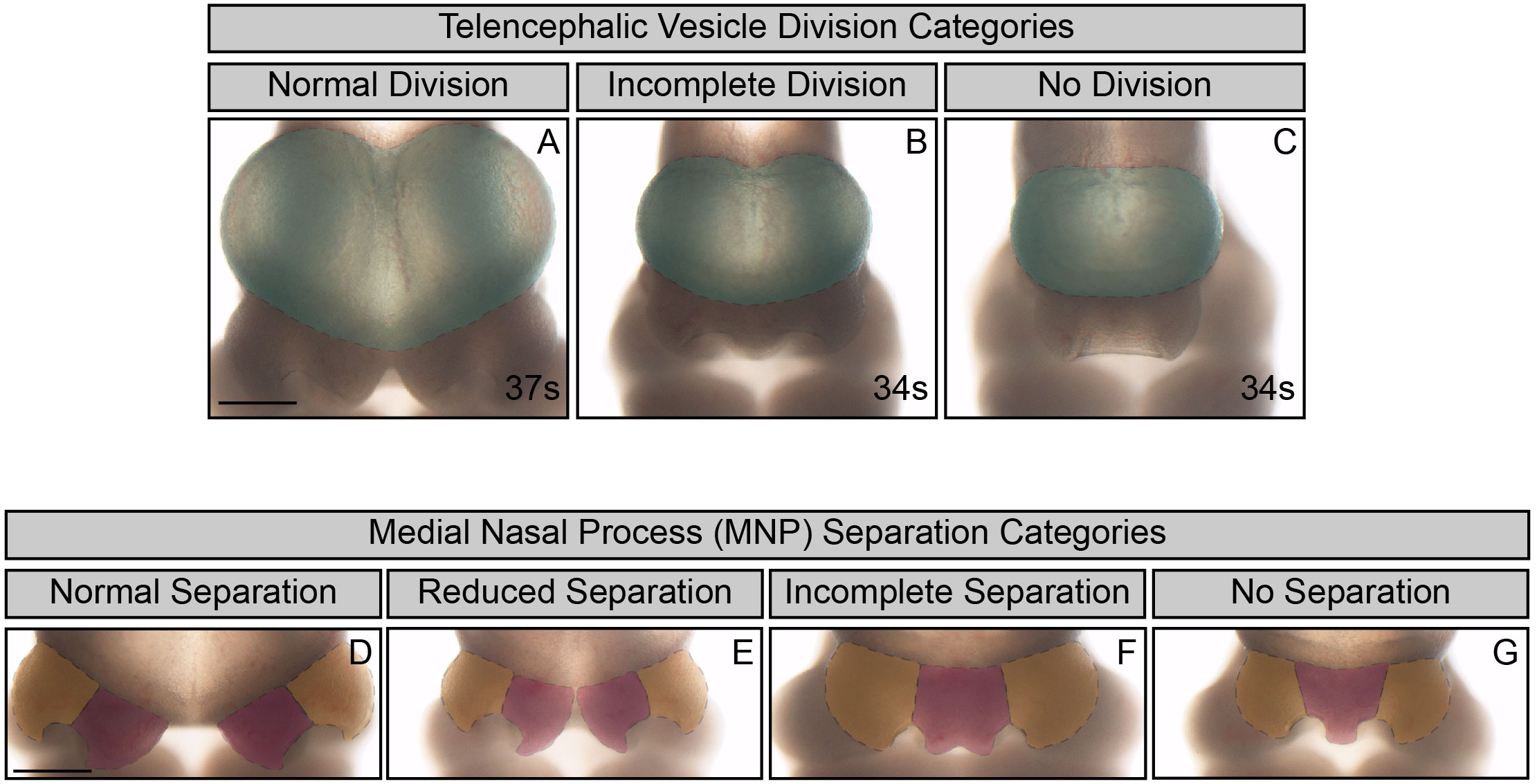
Definitions of categories used to quantify telencephalic vesicle division and MNP separation. En face view of E10.5 embryos (A-C). The telencephalic vesicles are pseudocolored in green and surrounded by a dotted line. Telencephalic vesicle division classification categories: normal division (A), incomplete division (B), no division (C). Midface view of E10.5 embryos (D-G). The lateral and medial nasal processes are pseudocolored in orange and red, respectively, and are surrounded by a dotted line. Medial nasal process (MNP) classification categories: normal separation (D), reduced separation (E), incomplete separation (F), and no separation (G). Scale bars (A, D), 500µm.

**Supplemental Figure 3.**
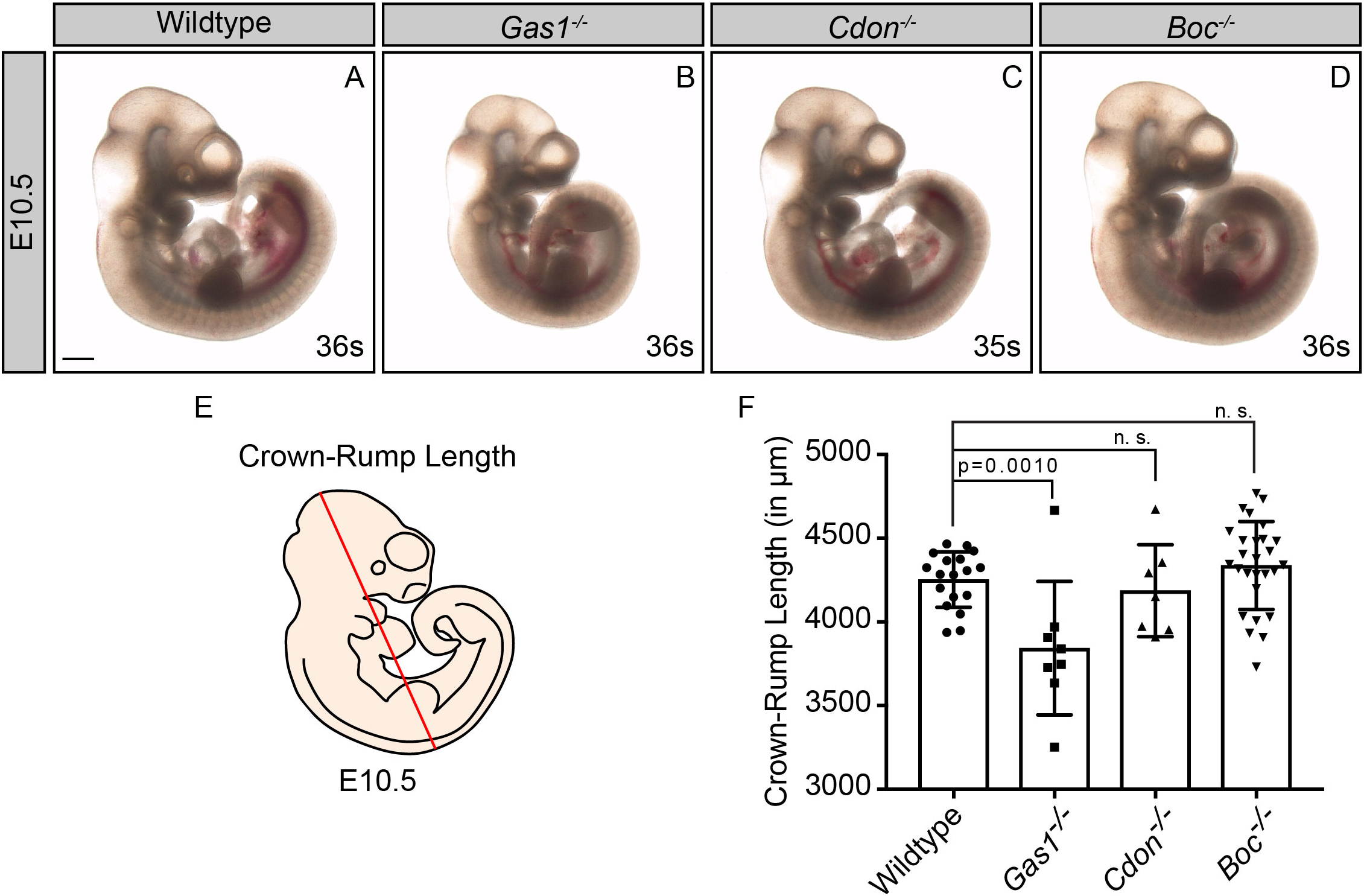
*Gas*1, but not *Cdon* or *Boc*, mutant embryos exhibit decreased embryo size at E10.5. Sagittal views of E10.5 embryos– wildtype (A), *Gas1^-/-^* (B), *Cdon^-/-^*(C), and *Boc^-/-^*(D). Schematic sagittal view of an E10.5 mouse embryo (E); the red diagonal line denotes crown-rump length. Crown-rump length quantitation in wildtype (n= 18), *Gas1^-/-^* (n=8), *Cdon^-/-^* (n=7), *Boc^-/-^* (n=27) embryos (F; in µm). Scale bar (A), 500 µm. Data are represented as the mean ± standard deviation. P-values were determined by a two-tailed Student’s *t*-test; n.s., not significant.

**Supplemental Figure 4.**
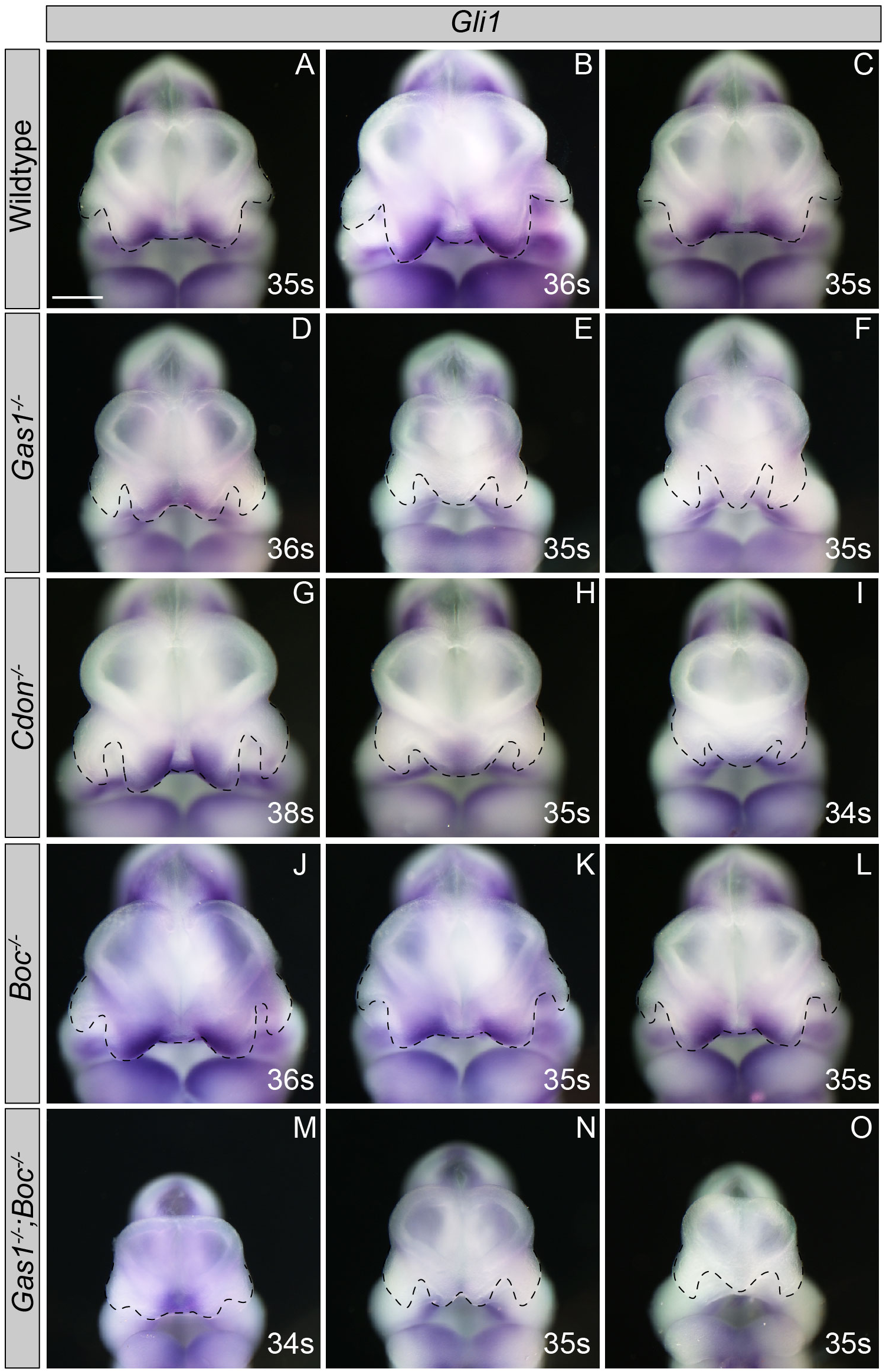
The Spectrum of HPE phenotypes correlates with changes in *Gli1* expression. *In situ* hybridization detection of *Gli1* expression in E10.5 forebrains (A-O). En face views of E10.5 forebrains– wildtype (A-C), *Gas1^-/-^* (D-F), *Cdon^-/-^* (G-I), *Boc^-/-^* (J-L) and *Gas1^-/-^;Boc^-/-^* embryos are shown. Somite number (s) is indicated in the lower right corner of each panel. Black dotted lines outline nasal processes. Notice that as the HPE phenotypes worsen (from left to right) in *Gas1* and *Cdon* mutants, the expression of *Gli1* in the MNP is lost. *Boc* mutants display equal levels of *Gli1* in the MNP and do not display any gross craniofacial defects. *Gas1;Boc* double mutants with rescue of the craniofacial defects (from left to right) maintain the expression of *Gli1* in the MNP, while mutants that do not display the rescue, the expression of *Gli1* is lost. Scale bars (A-O), 500 µm.

**Supplemental Figure 5.**
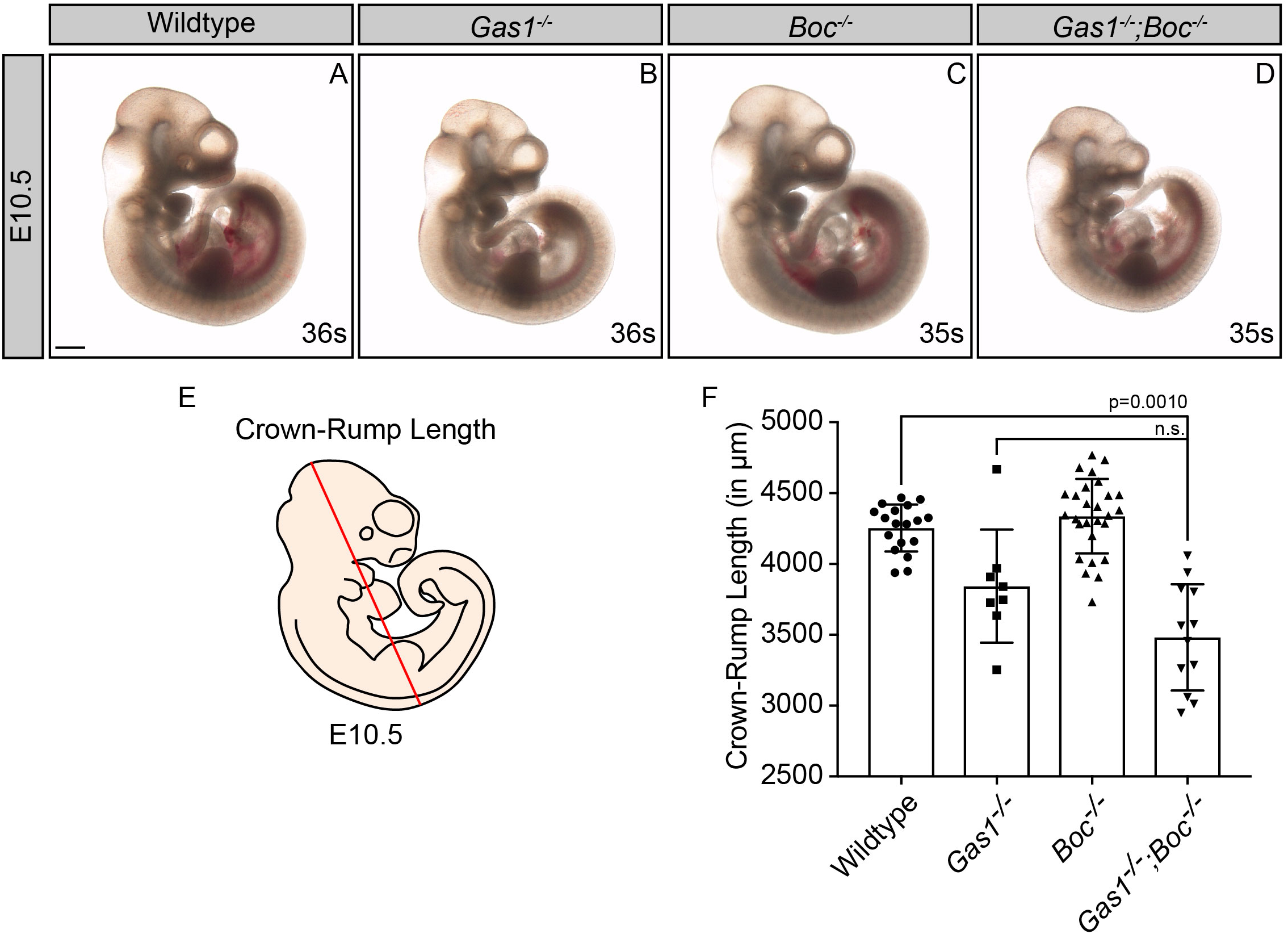
Reduced Crown-Rump Length in E10.5 *Gas1;Boc* double mutant embryos. Sagittal view of E10.5 wildtype (A), *Gas1^-/-^* (B), *Boc^-/-^* (C), and *Gas1^-/-^;Boc^-/-^* (D) embryos. Schematic sagittal view of an E10.5 mouse embryo; the red diagonal line denotes the crown-rump length (E). Crown-rump length quantitation (F; in µm). Scale bar in (A), 500µm. Data are represented as mean ± standard deviation. P-values were determined by two tailed Student’s *t*-test.

**Supplemental Figure 6.**
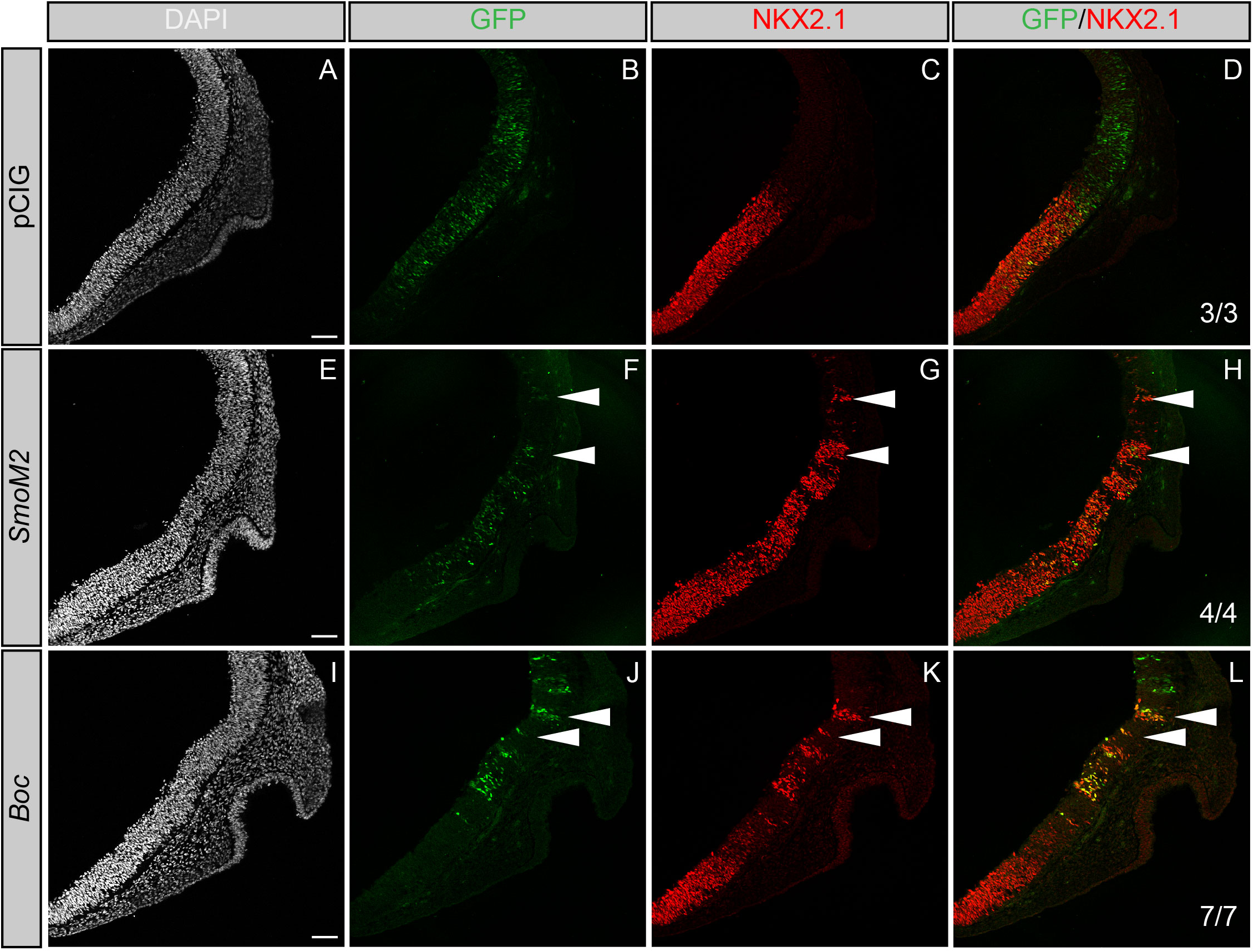
*Boc* promotes HH-dependent neural patterning in the developing chicken forebrain. Coronal sections of Hamburger-Hamilton stage 21-22 chicken telencephalons electroporated with empty vector (pCIG; A-D), *SmoM2* (E-H), and *Boc* (I-L). DAPI (grayscale; A,E,I) denotes nuclei. GFP+ cells (green; B,F,J) identify electroporated cells. Antibody detection of NKX2.1 (red; C,G,K) reads out HH pathway activity. Merged images are shown in (D,H,L). The number of electroporated embryos that display ectopic NKX2.1 expression is indicated in the lower right corner (D,H,L). White arrowheads highlight ectopic NKX2.1 expression. Scale bars in (A), (E), and (I), 50µm.

**Supplemental Figure 7.**
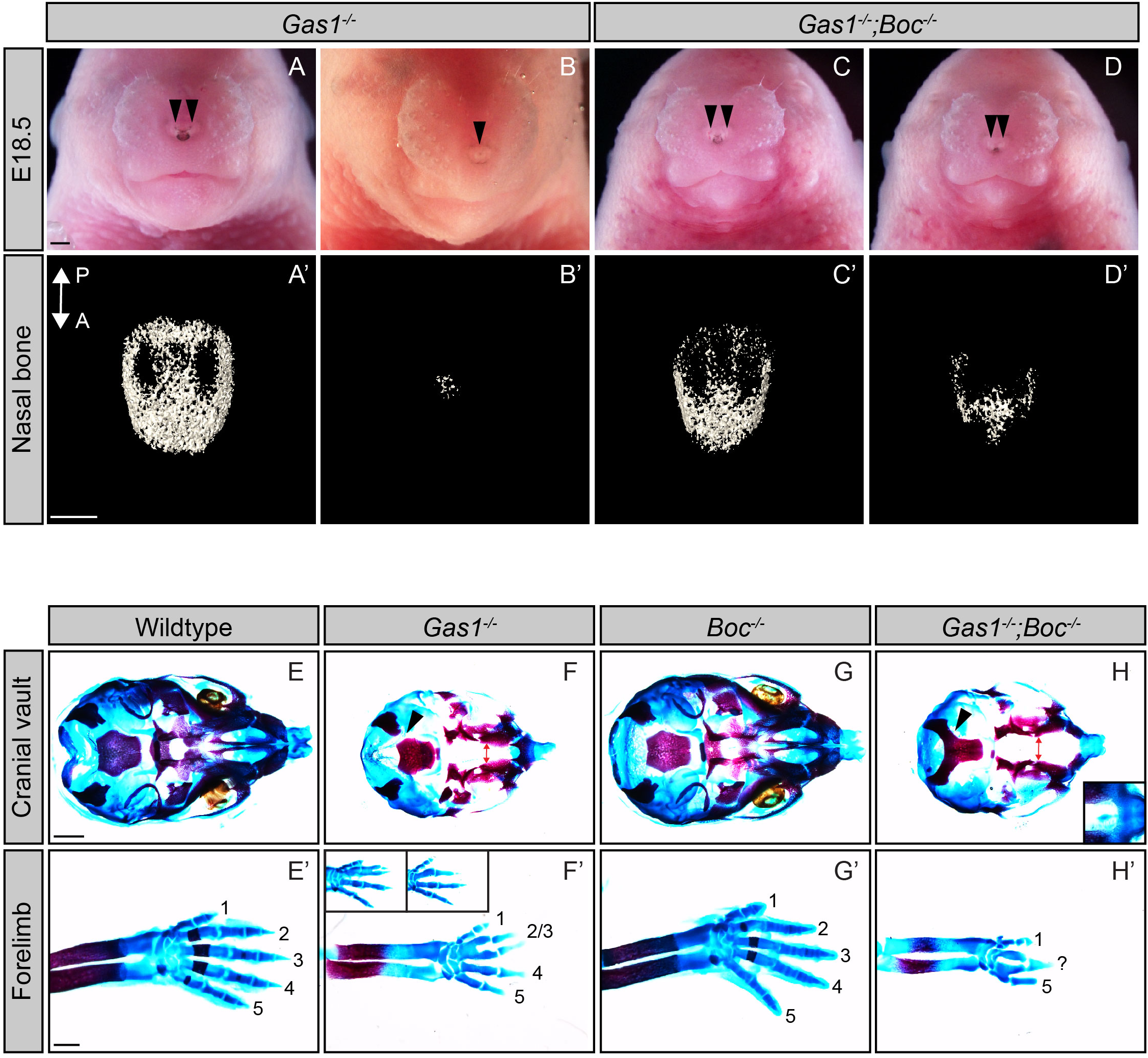
HPE phenotypes and digit specification defects in E18.5 *Gas1;Boc* mutant embryos. En face view of E18.5 *Gas1^-/-^* (A,B) and *Gas1^-/-^;Boc^-/-^* (C,D) embryos. Black arrowheads denote the nasal pits. Three dimensional reconstructions of microCT images of isolated nasal bones from E18.5 *Gas1^-/-^* (A’B’) and *Gas1^-/-^;Boc^-/-^* (C’,D’) embryos. A←→P specifies the anterior to posterior axis in (A’-D’). Ventral views of E18.5 cranial vaults from wildtype (E), *Gas1^-/-^* (F), *Boc^-/-^* (G), and *Gas1^-/-^;Boc^-/-^* (H) embryos, stained with alcian blue and alizarin red. Red double arrows denote the cleft palate in *Gas1^-/-^* and *Gas1^-/-^;Boc^-/-^* embryos and black arrowheads mark occipital bone. Inset in H indicates hypoplastic premaxilla in *Gas1^-/-^;Boc^-/-^* embryos. Forelimbs of E18.5 wildtype (E’), *Gas1^-/-^* (F’), *Boc^-/-^* (G’), and *Gas1^-/-^;Boc^-/-^* (H’) embryos, stained with alcian blue and alizarin red. Numbers denote specific digits where 1 is the most anterior and 5 is the most posterior. Insets in F demonstrate variable digit specification phenotypes in *Gas1^-/-^* embryos, which display either partial fusion of digits two and three (left), or the absence of either digit two or three (right). *Gas1^-/-^;Boc^-/-^* embryos exhibit a more severe limb phenotype where only digits 1 and 5 can be clearly identified; a third, unidentified digit is labeled with a question mark (Allen et al., 2011). Scale bars (A, A’, E, E’), 500 µm.

**Supplemental Figure 8.**
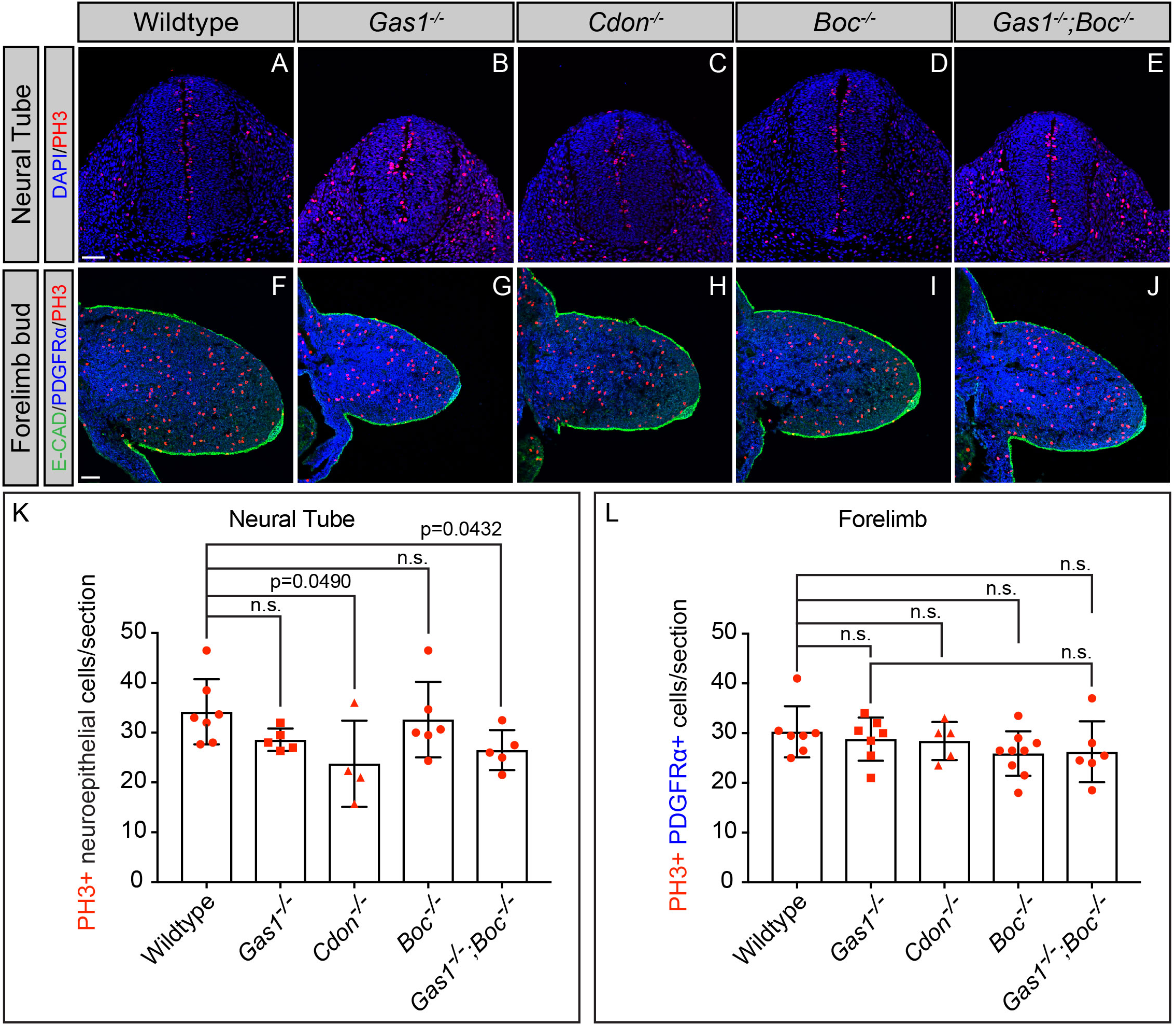
*Boc* does not contribute to neural tube or forelimb mesenchyme proliferation. Immunofluorescent analysis of proliferation in E10.5 neural tube (A-E) and forelimb (F-J) transverse sections from E10.5, wildtype (A,F), *Gas1^-/-^* (B,G), *Cdon^-/-^* (C,H), *Boc^-/-^* (D,I), and *Gas1^-/-^;Boc^-/-^* (E,J) embryos. Antibody detection of E-CADHERIN (E-CAD, green, F-J), PDGFRα (blue, F-J), and phospho-histone H3 (PH3, red, A-J). Nuclei are stained with DAPI (blue, A-E). Quantitation of PH3+ cells (2 sections/embryo) in the neural tube (K) and the forelimb bud (L) from E10.5 wildtype (n=6), *Gas1^-/-^* (n=5), *Boc^-/-^* (n=6) and *Gas1^-/-^;Boc^-/-^* (n=4) embryos. Data are presented as mean ± standard deviation. P-values were determined by two-tailed Student’s *t*-test. Scale bars (A,F), 50 µm.

**Supplemental table S1:**
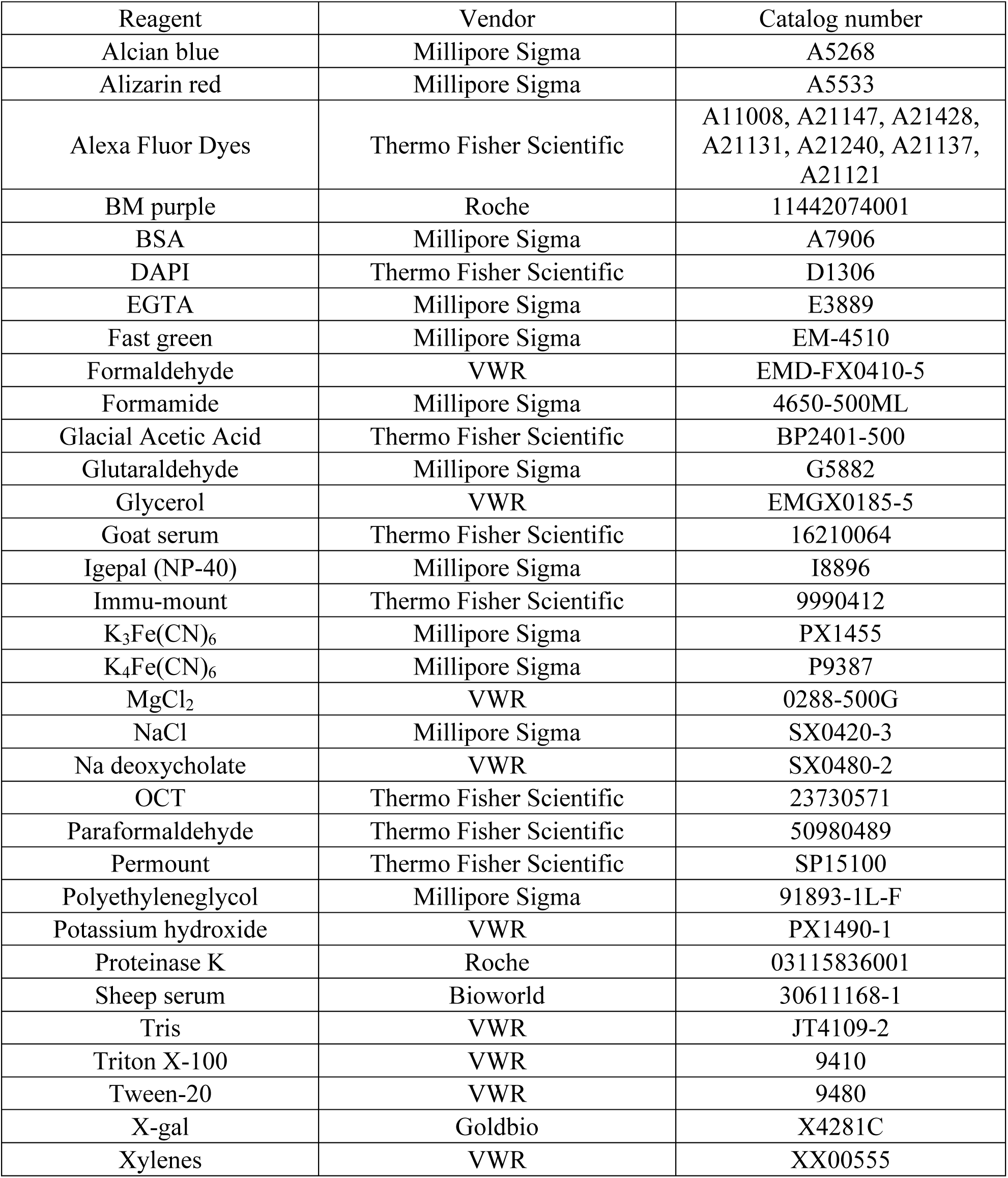
List of reagents

**Supplemental Table S2.** List of primary antibodies used for immunofluorescence

| Antibody | Vendor | Catalog number | Dilution |
| --- | --- | --- | --- |
| Digoxigenin | Roche | 11 093 274 910 | 1:4,000 |
| NKX2.1 (rabbit IgG) | Abcam | ab76013 | 1:200 |
| E-CADHERIN (mouseIgG2a) | BD-biosciences | 610181 | 1:500 |
| NKX2.2 (mouseIgG2b) | Developmental Studies Hybridoma Bank | 74.5A5 | 1:20 |
| OLIG2 (rabbit IgG) | Millipore Sigma | AB9610 | 1:2,000 |
| NKX6.1 (mouseIgG1) | Developmental Studies Hybridoma Bank | F55A10 | 1:20 |
| Phospho-histone H3 (rabbit IgG) | Millipore Sigma | 06-570 | 1:1,000 |
| Phospho-histone H3 (mouse IgG1) | Cell Signaling | 9706S | 1:100 |
| PDGFRα (rabbit IgG) | Cell Signaling | 3174S | 1:100 |

## References

Allen, B. L., Song, J. Y., Izzi, L., Althaus, I. W., Kang, J. S., Charron, F., Krauss, R. S. and McMahon, A. P. (2011). Overlapping roles and collective requirement for the coreceptors GAS1, CDO, and BOC in SHH pathway function. Dev Cell 20, 775–787.

Allen, B. L., Tenzen, T. and McMahon, A. P. (2007). The Hedgehog-binding proteins Gas1 and Cdo cooperate to positively regulate Shh signaling during mouse development. Genes Dev 21, 1244–1257.

Aoto, K., Shikata, Y., Imai, H., Matsumaru, D., Tokunaga, T., Shioda, S., Yamada, G. and Motoyama, J. (2009). Mouse Shh is required for prechordal plate maintenance during brain and craniofacial morphogenesis. Dev Biol 327, 106–120.

Bae, G. U., Domene, S., Roessler, E., Schachter, K., Kang, J. S., Muenke, M. and Krauss, R. S. (2011). Mutations in CDON, encoding a hedgehog receptor, result in holoprosencephaly and defective interactions with other hedgehog receptors. Am J Hum Genet 89, 231–240.

Beachy, P. A., Hymowitz, S. G., Lazarus, R. A., Leahy, D. J. and Siebold, C. (2010). Interactions between Hedgehog proteins and their binding partners come into view. Genes Dev 24, 2001–2012.

Bergeron, S. A., Tyurina, O. V., Miller, E., Bagas, A. and Karlstrom, R. O. (2011). Brother of cdo (umleitung) is cell-autonomously required for Hedgehog-mediated ventral CNS patterning in the zebrafish. Development 138, 75–85.

Briscoe, J. and Therond, P. P. (2013). The mechanisms of Hedgehog signalling and its roles in development and disease. Nat Rev Mol Cell Biol 14, 416–429.

Brugmann, S. A., Allen, N. C., James, A. W., Mekonnen, Z., Madan, E. and Helms, J. A. (2010). A primary cilia-dependent etiology for midline facial disorders. Hum Mol Genet 19, 1577–1592.

Cabrera, J. R., Sanchez-Pulido, L., Rojas, A. M., Valencia, A., Manes, S., Naranjo, J. R. and Mellstrom, B. (2006). Gas1 is related to the glial cell-derived neurotrophic factor family receptors alpha and regulates Ret signaling. J Biol Chem 281, 14330–14339.

Cardozo, M. J., Sanchez-Arrones, L., Sandonis, A., Sanchez-Camacho, C., Gestri, G., Wilson, S. W., Guerrero, I. and Bovolenta, P. (2014). Cdon acts as a Hedgehog decoy receptor during proximal-distal patterning of the optic vesicle. Nat Commun 5, 4272.

Chiang, C., Litingtung, Y., Lee, E., Young, K. E., Corden, J. L., Westphal, H. and Beachy, P. A. (1996). Cyclopia and defective axial patterning in mice lacking Sonic hedgehog gene function. Nature 383, 407–413.

Cobourne, M. T., Miletich, I. and Sharpe, P. T. (2004). Restriction of sonic hedgehog signalling during early tooth development. Development 131, 2875–2885.

Cole, F. and Krauss, R. S. (2003). Microform holoprosencephaly in mice that lack the Ig superfamily member Cdon. Curr Biol 13, 411–415.

Cordero, D., Marcucio, R., Hu, D., Gaffield, W., Tapadia, M. and Helms, J. A. (2004). Temporal perturbations in sonic hedgehog signaling elicit the spectrum of holoprosencephaly phenotypes. J Clin Invest 114, 485–494.

Dai, P., Akimaru, H., Tanaka, Y., Maekawa, T., Nakafuku, M. and Ishii, S. (1999). Sonic Hedgehog-induced activation of the Gli1 promoter is mediated by GLI3. J Biol Chem 274, 8143–8152.

Dessaud, E., McMahon, A. P. and Briscoe, J. (2008). Pattern formation in the vertebrate neural tube: a sonic hedgehog morphogen-regulated transcriptional network. Development 135, 2489–2503.

Ho, T. V., Iwata, J., Ho, H. A., Grimes, W. C., Park, S., Sanchez-Lara, P. A. and Chai, Y. (2015). Integration of comprehensive 3D microCT and signaling analysis reveals differential regulatory mechanisms of craniofacial bone development. Dev Biol 400, 180–190.

Holtz, A. M., Peterson, K. A., Nishi, Y., Morin, S., Song, J. Y., Charron, F., McMahon, A. P. and Allen, B. L. (2013). Essential role for ligand-dependent feedback antagonism of vertebrate hedgehog signaling by PTCH1, PTCH2 and HHIP1 during neural patterning. Development 140, 3423–3434.

Hong, M. and Krauss, R. S. (2018). Modeling the complex etiology of holoprosencephaly in mice. Am J Med Genet C Semin Med Genet 178, 140–150.

Hong, M., Srivastava, K., Kim, S., Allen, B. L., Leahy, D. J., Hu, P., Roessler, E., Krauss, R. S. and Muenke, M. (2017). BOC is a modifier gene in holoprosencephaly. Hum Mutat 38, 1464–1470.

Hu, D. and Helms, J. A. (1999). The role of sonic hedgehog in normal and abnormal craniofacial morphogenesis. Development 126, 4873–4884.

Hui, C. C. and Angers, S. (2011). Gli proteins in development and disease. Annu Rev Cell Dev Biol 27, 513–537.

Izzi, L., Levesque, M., Morin, S., Laniel, D., Wilkes, B. C., Mille, F., Krauss, R. S., McMahon, A. P., Allen, B. L. and Charron, F. (2011). Boc and Gas1 each form distinct Shh receptor complexes with Ptch1 and are required for Shh-mediated cell proliferation. Dev Cell 20, 788–801.

Jeong, J., Mao, J., Tenzen, T., Kottmann, A. H. and McMahon, A. P. (2004). Hedgehog signaling in the neural crest cells regulates the patterning and growth of facial primordia. Genes Dev 18, 937–951.

Jiang, R., Bush, J. O. and Lidral, A. C. (2006). Development of the upper lip: morphogenetic and molecular mechanisms. Dev Dyn 235, 1152–1166.

Kang, J. S., Gao, M., Feinleib, J. L., Cotter, P. D., Guadagno, S. N. and Krauss, R. S. (1997). CDO: an oncogene-, serum-, and anchorage-regulated member of the Ig/fibronectin type III repeat family. J Cell Biol 138, 203–213.

Kang, J. S., Mulieri, P. J., Hu, Y., Taliana, L. and Krauss, R. S. (2002). BOC, an Ig superfamily member, associates with CDO to positively regulate myogenic differentiation. EMBO J 21, 114–124.

Krauss, R. S. (2007). Holoprosencephaly: new models, new insights. Expert Rev Mol Med 9, 1–17.

Kuwajima, T., Sitko, A. A., Bhansali, P., Jurgens, C., Guido, W. and Mason, C. (2013). ClearT: a detergent- and solvent-free clearing method for neuronal and non-neuronal tissue. Development 140, 1364–1368.

Lee, C. S., Buttitta, L. and Fan, C. M. (2001). Evidence that the WNT-inducible growth arrest-specific gene 1 encodes an antagonist of sonic hedgehog signaling in the somite. Proc Natl Acad Sci U S A 98, 11347–11352.

Lei, Q., Jeong, Y., Misra, K., Li, S., Zelman, A. K., Epstein, D. J. and Matise, M. P. (2006). Wnt signaling inhibitors regulate the transcriptional response to morphogenetic Shh-Gli signaling in the neural tube. Dev Cell 11, 325–337.

Lum, L., Yao, S., Mozer, B., Rovescalli, A., Von Kessler, D., Nirenberg, M. and Beachy, P. A. (2003). Identification of Hedgehog pathway components by RNAi in Drosophila cultured cells. Science 299, 2039–2045.

Makihara, S., Morin, S., Ferent, J., Cote, J. F., Yam, P. T. and Charron, F. (2018). Polarized Dock Activity Drives Shh-Mediated Axon Guidance. Dev Cell 46, 410–425 e417.

Marcucio, R. S., Cordero, D. R., Hu, D. and Helms, J. A. (2005). Molecular interactions coordinating the development of the forebrain and face. Dev Biol 284, 48–61.

Marigo, V., Davey, R. A., Zuo, Y., Cunningham, J. M. and Tabin, C. J. (1996). Biochemical evidence that patched is the Hedgehog receptor. Nature 384, 176–179.

Martinelli, D. C. and Fan, C. M. (2007). Gas1 extends the range of Hedgehog action by facilitating its signaling. Genes Dev 21, 1231–1243.

McLellan, J. S., Zheng, X., Hauk, G., Ghirlando, R., Beachy, P. A. and Leahy, D. J. (2008). The mode of Hedgehog binding to Ihog homologues is not conserved across different phyla. Nature 455, 979–983.

McMahon, A. P., Ingham, P. W. and Tabin, C. J. (2003). Developmental roles and clinical significance of hedgehog signaling. Curr Top Dev Biol 53, 1–114.

Muenke, M. and Beachy, P. A. (2000). Genetics of ventral forebrain development and holoprosencephaly. Curr Opin Genet Dev 10, 262–269.

Ohazama, A., Haycraft, C. J., Seppala, M., Blackburn, J., Ghafoor, S., Cobourne, M., Martinelli, D. C., Fan, C. M., Peterkova, R., Lesot, H., et al. (2009). Primary cilia regulate Shh activity in the control of molar tooth number. Development 136, 897–903.

Okada, A., Charron, F., Morin, S., Shin, D. S., Wong, K., Fabre, P. J., Tessier-Lavigne, M. and McConnell, S. K. (2006). Boc is a receptor for sonic hedgehog in the guidance of commissural axons. Nature 444, 369–373.

Pabst, O., Herbrand, H., Takuma, N. and Arnold, H. H. (2000). NKX2 gene expression in neuroectoderm but not in mesendodermally derived structures depends on sonic hedgehog in mouse embryos. Dev Genes Evol 210, 47–50.

Ribeiro, L. A., Quiezi, R. G., Nascimento, A., Bertolacini, C. P. and Richieri-Costa, A. (2010). Holoprosencephaly and holoprosencephaly-like phenotype and GAS1 DNA sequence changes: Report of four Brazilian patients. Am J Med Genet A 152A, 1688–1694.

Roessler, E., El-Jaick, K. B., Dubourg, C., Velez, J. I., Solomon, B. D., Pineda-Alvarez, D. E., Lacbawan, F., Zhou, N., Ouspenskaia, M., Paulussen, A., et al. (2009). The mutational spectrum of holoprosencephaly-associated changes within the SHH gene in humans predicts loss-of-function through either key structural alterations of the ligand or its altered synthesis. Hum Mutat 30, E921–935.

Roessler, E. and Muenke, M. (2010). The molecular genetics of holoprosencephaly. Am J Med Genet C Semin Med Genet 154C, 52–61.

Rubenstein, J. L. and Beachy, P. A. (1998). Patterning of the embryonic forebrain. Curr Opin Neurobiol 8, 18–26.

Schachter, K. A. and Krauss, R. S. (2008). Murine models of holoprosencephaly. In Curr Top Dev Biol, pp. 139–170.

Schneider, C. A., Rasband, W. S. and Eliceiri, K. W. (2012). NIH Image to ImageJ: 25 years of image analysis. Nat Methods 9, 671–675.

Seppala, M., Depew, M. J., Martinelli, D. C., Fan, C. M., Sharpe, P. T. and Cobourne, M. T. (2007). Gas1 is a modifier for holoprosencephaly and genetically interacts with sonic hedgehog. J Clin Invest 117, 1575–1584.

Seppala, M., Xavier, G. M., Fan, C. M. and Cobourne, M. T. (2014). Boc modifies the spectrum of holoprosencephaly in the absence of Gas1 function. Biol Open 3, 728–740.

Tenzen, T., Allen, B. L., Cole, F., Kang, J. S., Krauss, R. S. and McMahon, A. P. (2006). The cell surface membrane proteins Cdo and Boc are components and targets of the Hedgehog signaling pathway and feedback network in mice. Dev Cell 10, 647–656.

Vuong, T. A., Leem, Y. E., Kim, B. G., Cho, H., Lee, S. J., Bae, G. U. and Kang, J. S. (2017). A Sonic hedgehog coreceptor, BOC regulates neuronal differentiation and neurite outgrowth via interaction with ABL and JNK activation. Cell Signal 30, 30–40.

Wang, H., Lei, Q., Oosterveen, T., Ericson, J. and Matise, M. P. (2011). Tcf/Lef repressors differentially regulate Shh-Gli target gene activation thresholds to generate progenitor patterning in the developing CNS. Development 138, 3711–3721.

Wilkinson, D. G. (1992). In situ hybridization : a practical approach. Oxford ; New York: IRL Press at Oxford University Press.

Xavier, G. M., Seppala, M., Barrell, W., Birjandi, A. A., Geoghegan, F. and Cobourne, M. T. (2016a). Hedgehog receptor function during craniofacial development. Dev Biol 415, 198–215.

Xavier, G. M., Seppala, M., Papageorgiou, S. N., Fan, C. M. and Cobourne, M. T. (2016b). Genetic interactions between the hedgehog co-receptors Gas1 and Boc regulate cell proliferation during murine palatogenesis. Oncotarget 7, 79233–79246.

Xie, J., Murone, M., Luoh, S. M., Ryan, A., Gu, Q., Zhang, C., Bonifas, J. M., Lam, C. W., Hynes, M., Goddard, A., et al. (1998). Activating Smoothened mutations in sporadic basal-cell carcinoma. Nature 391, 90–92.

Xu, J., Liu, H., Lan, Y., Adam, M., Clouthier, D. E., Potter, S. and Jiang, R. (2019). Hedgehog signaling patterns the oral-aboral axis of the mandibular arch. Elife 8.

Yao, S., Lum, L. and Beachy, P. (2006). The ihog cell-surface proteins bind Hedgehog and mediate pathway activation. Cell 125, 343–357.

Zhang, W., Hong, M., Bae, G. U., Kang, J. S. and Krauss, R. S. (2011). Boc modifies the holoprosencephaly spectrum of Cdo mutant mice. Dis Model Mech 4, 368–380.

Zhang, W., Kang, J. S., Cole, F., Yi, M. J. and Krauss, R. S. (2006). Cdo functions at multiple points in the Sonic Hedgehog pathway, and Cdo-deficient mice accurately model human holoprosencephaly. Dev Cell 10, 657–665.

